# Transcriptome profile of the zebrafish atrioventricular canal reveals molecular signatures of pacemaker and valve mesenchyme

**DOI:** 10.1101/2021.01.27.428386

**Authors:** Abu Nahia Karim, Migdał Maciej, Quinn T. Alexander, Poon Kar-Lai, Łapinski Maciej, Sulej Agata, Pawlak Michał, Bugajski Łukasz, Piwocka Katarzyna, Brand Thomas, Kohl Peter, Korzh Vladimir, Winata Cecilia

## Abstract

The atrioventricular canal (AVC) is an essential feature of the heart, which separates the atrium from the ventricle. During heart morphogenesis, it is a hub of molecular processes necessary for distinguishing heart regions; most importantly, for the formation of the AV conduction system and cardiac valves. To better understand the molecular processes underlying AVC development and function, we utilized the transgenic zebrafish line *sqet31Et* with EGFP expression in the AVC region to isolate this cell population by FACS and profiled its transcriptome by RNA-seq at 48 and 72 hours post fertilization (hpf). Compared to the rest of the heart, the AVC is enriched for the expression of molecular markers associated with mammalian AVC and AV node, including *cx36.7* and *cx45* which encode connexins forming low conductance gap junctions. Using the transgenic line *Tg(myl7:mermaid)* encoding the voltage-sensitive fluorescent protein, we showed that loss of function of Isl1 abolished the pacemaker-containing sinoatrial ring (SAR) and resulted in an erratic spread of excitation pattern from the SAR to AVC, indicating the dysfunction of the primary pacemaker. Concurrently, ectopic excitation in the AVC region was observed, suggesting that the zebrafish AVC possesses inherent automaticity although insufficient to replace the primary pacemaking activity of the SAR. Comparisons between the SAR and AVC transcriptomes revealed partially overlapping expression profiles of various ion channels and gap junction proteins which reflects their diversified functions. Lastly, we observed dynamic expression of epithelial-to-mesenchymal transition markers, as well as components of TGF-β, Notch, and Wnt signaling pathways, which have been implicated in the formation of AVC conduction and cardiac valves. Our results uncovered the molecular hallmarks of the developing AVC region and demonstrated its role in the structural and electrophysiological separation between the atrium and ventricle.

**Author summary:** The atrioventricular canal is a structure in the embryonic heart which separates the atrium from the ventricle. It gives rise to the AV node and cardiac valves - two important structures which ensure unidirectional blood flow between heart chambers. The AV node serves to regulate the propagation of electrical impulses between the two chambers, such that they contract consecutively. Using the zebrafish as model organism, we performed gene expression profiling and characterized electrical conduction patterns between the sinoatrial primary pacemaker and AVC. We discovered that the zebrafish AVC possesses similar features to the mammalian AV node, including slow conduction, inherent pacemaking activity, and the expression of conserved developmental genes. The molecular profile of the AVC is distinct from that of the sinoatrial pacemaker, which reflects their distinct roles. In addition, we found that genes regulating cardiac valve development were also expressed in the AVC, illustrating the importance of this region for establishing both electrophysiological and structural separation between the heart chambers. Besides establishing conserved aspects between zebrafish and mammalian conduction system, the data generated in this study constitutes a valuable resource for studying AVC development and discovery of novel candidate genes implicated in regulating cardiac rhythm and cardiac valve formation.

## Introduction

The atrioventricular canal (AVC) serves two critical purposes in cardiac development and function. Firstly, the AVC gives rise to the AV node, which constitutes part of the cardiac conduction system (CCS) responsible for generating and transmitting electrical impulses necessary for coordinated heart contraction [1, 2]. In the mammalian heart, the AV node can be found within the interatrial septum, at the AV junction [3]. The AV node constitutes a group of cells regulating the transmission of electrical impulses between cardiac chambers. Electrical impulses originating from the sinoatrial (SA) node are delayed by a fraction of a second in the AV node before being propagated to the ventricle. Such delay ensures consecutive contractions of the atrium and ventricle [4]. In the mature, 4-chambered murine heart, the AV node serves as the only region of continuity between atrial and ventricular myocardium. It is surrounded by low-conducting fibrous tissue known as the annulus fibrosus, which derives from the epicardium and acts as an insulator between the atrial and ventricular myocardium [5]. The AV node is also often referred to as a secondary pacemaker as it possesses intrinsic automaticity, which renders it a potential arrhythmogenic source in cases where weakened or abnormal impulses from the SA node are not able to override it [3,6,7].

In the mammalian embryonic heart, the AV myocardium is slow conducting, unlike chamber myocardium. This property is retained in those cells that become the future AV node in the adult heart [2]. These cells express *Bmp4, Tbx2,* and *Tbx3,* which suppress the genetic program leading to the specification of working cardiomyocytes [8–10]. The electrophysiological properties of the AV node are determined by several factors, mainly the electrical coupling between its cells mediated by gap junctions. Connexins form gap junctions by either homogenous or heterogeneous pairings, resulting in a different range of conductivity [11, 12]. In the mammalian heart, CX30.2 and CX45 form low or ultra-low conductance gap junctions and are enriched in AV pacemaker cells [13, 14]. CX43 typically forms medium conductance gap junctions in the working myocardium [15, 16]. It can also form low-conductance gap junctions by pairing with lower conduction connexins such as CX30.2 [17].

In zebrafish, the earliest indication of a functional AV-node-like region in the heart was observed through calcium wave imaging, which revealed significant conductance delay between the atrium and ventricle from 36 hpf [18]. Notch1b and Neuregulin expressed in the endocardium have been shown to be involved in the development of this conduction delay [19]. To date, no live markers of the pacemaker have been available for direct visualization of its morphology and structure, or isolation for molecular characterization. Zebrafish orthologs of *Tbx2,* and *Tbx3* are expressed in the region that corresponds to the AVC in the mammalian heart [20, 21]; however, no detailed analyses have been made to elucidate the molecular profile of this structure, nor confirm its homologous function to the mammalian AV node. The LIM-homeodomain transcription factor (TF) Islet1 (Isl1) [22–24] was found to play a role in the development of the primary pacemaker, the SA node, as its deficiency causes cardiac arrhythmia [21, 25]. Isl1-positive cells in the sinoatrial region (SAR) of adult zebrafish co-expresses *hcn4*, which encodes the hyperpolarization-activated channel responsible for generating the pacemaker current (If) [26]. In the AVC of adult zebrafish, a small group of *hcn4*-positive cells was found in the AV valves. However, in contrast to the SAR, Hcn4-positive cells in the AVC region were Isl1-negative [27]. The earliest expression of *hcn4* in zebrafish embryonic AVC was reported from 52 hpf [28], which suggests that it could potentially function as a secondary pacemaker. Electrical silencing of cells in the SAR region of the embryonic zebrafish heart using optogenetics abolished the heartbeat, which suggests that the activities of alternative pacemaker regions, such as the AVC, are not sufficient to drive heart contractions [29]. Interestingly, surgical isolation of the ventricle from the atrium led to the establishment of the AV region as the site of electrical activation origin, which revealed its ability to function as a secondary pacemaker although with a slower excitation rate [27].

Besides its role in cardiac conduction, the AVC gives rise to the atrial and ventricular septa, which provide structural division between the four chambers of the mammalian heart. These structures also include the primary cardiac valves: the mitral and tricuspid valves. Studies in various model organisms have revealed the stepwise processes in AVC development, including its early patterning, endocardial cushion formation, and valve maturation [2, 30]. During cardiac chamber formation, a layer of extracellular matrix known as the cardiac jelly separates the myocardium from the endocardium. The cardiac jelly subsequently disappears from the chamber-forming regions, while being retained at the AVC region. Signals from the myocardium induce the endocardial cells to undergo epithelial-to-mesenchymal-transition (EMT), a process where epithelial cells undergo changes in cell polarity and cell-cell adhesion properties and are converted to migratory mesenchymal cells [31]. These cells populate the underlying cardiac jelly, becoming the mesenchyme substrate [32], which forms the endocardial cushions. These cushions continue to grow, forming the septa separating ventricles and atria, as well as valve leaflets. TGF-β signaling is known to play a role in endocardial cushion formation by promoting EMT [33]. In addition, Notch, Wnt, and BMP signaling activities in AVC endocardium and myocardium are also crucial to regulate the formation of endocardial cushion and valve [34].

In the zebrafish heart, AVC formation is initiated as early as 30 hpf when a constriction between the atrium and ventricle separates the two chambers as the heart begins to loop. At the same time, endocardial cells at the A-V border start becoming cuboidal in shape, and cardiomyocytes in the same region adopt a trapezoid morphology with a wider basolateral surface. These cells express higher levels of the surface adhesion molecule *alcama* compared to working cardiomyocytes [35]. Around the time when heart looping is initiated at 36 hpf, *bmp4* expression becomes restricted to the AVC myocardium [36], where it plays a role in the formation of cardiac jelly together with Has2 [37]. The development of valves is initiated by the formation of endocardial cushions at two opposite sides of the AVC at 55 hpf [35]. By 60 hpf, some of these endocardial cells migrate into the underlying cardiac jelly and undergo EMT, which provides the substrate for valve development [35]. Canonical Wnt signaling is required for zebrafish endocardial cushion formation [38, 39], although its precise mechanism is still unknown. In addition, Wnt signals originating from the endocardium induce myocardial *bmp4* and *tbx2b* expression, which are necessary for patterning the AVC myocardium [40]. Similarly, *notch1b* and its ligand *dll4* are expressed in the AVC endocardium [34, 36] and are required for the formation of the endocardial cushion as well as AV conduction tissue [35, 41]. Given its important role in the formation of major cardiac structures, disruptions to AVC development can result in various forms of septal defects, as well as valve abnormalities leading to heart failure. In addition, defects of the AV node may lead to various degrees of AV block, which gives rise to cardiac arrhythmia [42]. However, despite its importance, our knowledge of genetic events responsible for the development and function of the AVC region in general, as well as of the AV node in particular, is still limited. One of the main challenges in studying the AVC is posed by their complex spatial anatomy and cellular heterogeneity [43]. For this reason, isolation of specific cell populations from ambient working myocardium and other surrounding tissues is challenging, which limits the identification of clear-cut molecular markers. Moreover, functional genetic studies of the AVC in higher vertebrates are impractical due to the early embryonic lethality caused by loss of function of essential genes. Thus, there is still a lack of a good and established system that can reliably model the development, physiology, and pathology of the AVC.

Despite the significant evolutionary distance between human and zebrafish, the zebrafish heart is remarkably similar to the human heart in terms of basal heart rate, electrophysiological properties, and action potential shape and duration [44, 45]. Several zebrafish mutants for cardiac ion channels have been described, which display phenotypes closely resembling those found in various forms of human arrhythmia [46–48]. In addition, relevant phenotypes have also been shown to result from transgenic expression of human disease mutations, which illustrates the high conservation of molecular pathways regulating electrical conduction in the heart [49]. The zebrafish therefore holds great potential to model human pathologies affecting the AVC and AV pacemaker function.

An enhancer trap screen performed in zebrafish has generated a collection of transgenic lines expressing enhanced green fluorescent protein (EGFP) in different tissues or subdomains of the heart [44, 50]. Among these lines, two express EGFP either in the SAR or AVC. The transgenic line *sqet33mi59BEt*, in which the enhancer trap was inserted close by the *fhf2* gene locus, expresses EGFP in the SAR [51]. Recently, we completed the analysis of the transcriptome of these cells [52]. The transgenic line *sqet31Et* exhibits green fluorescent protein (GFP) expression in a ring structure in the region of the AVC [50], which corresponds to the ring of AV conduction tissue as described previously [18, 19]. Here we utilize the *in vivo* labeling of the AVC in the transgenic line *sqet31Et* to isolate cells making up this structure and perform detailed molecular characterization by transcriptome profiling at 48 hpf and 72 hpf, corresponding to the time of CCS and cardiac valve development. To better understand the physiology of the CCS in zebrafish, we characterized the electrical conduction patterns between the SAR and AVC, and cross-compared the transcriptome profiles of both pacemaker regions. Our results show that the AVC gene expression profile exhibits hallmarks of the mammalian AV node and reflects ongoing biological processes implicated in valve development. Comparisons between the SAR and AVC transcriptomes revealed differences reflected in expression profiles of ion channels and connexins implicated in pacemaker function. This data constitutes a valuable resource for the study of AVC development and function and identification of candidate genes implicated in these processes.

## Results

### Transgenic zebrafish line *sqet31Et* expresses EGFP in the zebrafish AVC

The *sqet31Et* transgenic line expresses EGFP in a ring structure marking the AVC, which likely corresponds to slow conducting myocytes homologous to the mammalian AV node [18,19,50,51]. To better visualize this structure, we performed high resolution imaging at 48 hpf and 72 hpf (Fig. 1A-D). To mark the myocardium, we crossed *sqet31Et* with *Tg(myl7:mRFP)* that expresses membrane-bound RFP (mRFP) in cardiomyocytes. Confocal imaging of the AVC region revealed that at the surface of the AVC, the EGFP and mRFP expression overlapped, confirming the myocardial nature of the EGFP-expressing cells (Fig. 1C). At 72 hpf, two additional groups of ∼3 cuboidal-shaped cells were detected at the deeper layer facing the cardiac lumen (Fig. 1D). These cells appear to be a part of the characteristic protrusion into the cardiac lumen most likely representing the developing atrioventricular cushion. Based on their location between the endocardium and myocardium, these cells have been previously defined as constituents of the non-chamber valve tissue [50, 51]. An additional EGFP expression domain detected previously [50] was observed at the bulbus arteriosus (BA, corresponding to the mesenchyme of the outflow tract in mammals [53]) from 72 hpf (Fig. 1B). The EGFP expressing domain in the developing heart of the *sqet31Et* transgenic line thus consists of largely AVC myocardium, with another expression domain in the BA, the latter observed only at 72 hpf.

**Figure 1.**
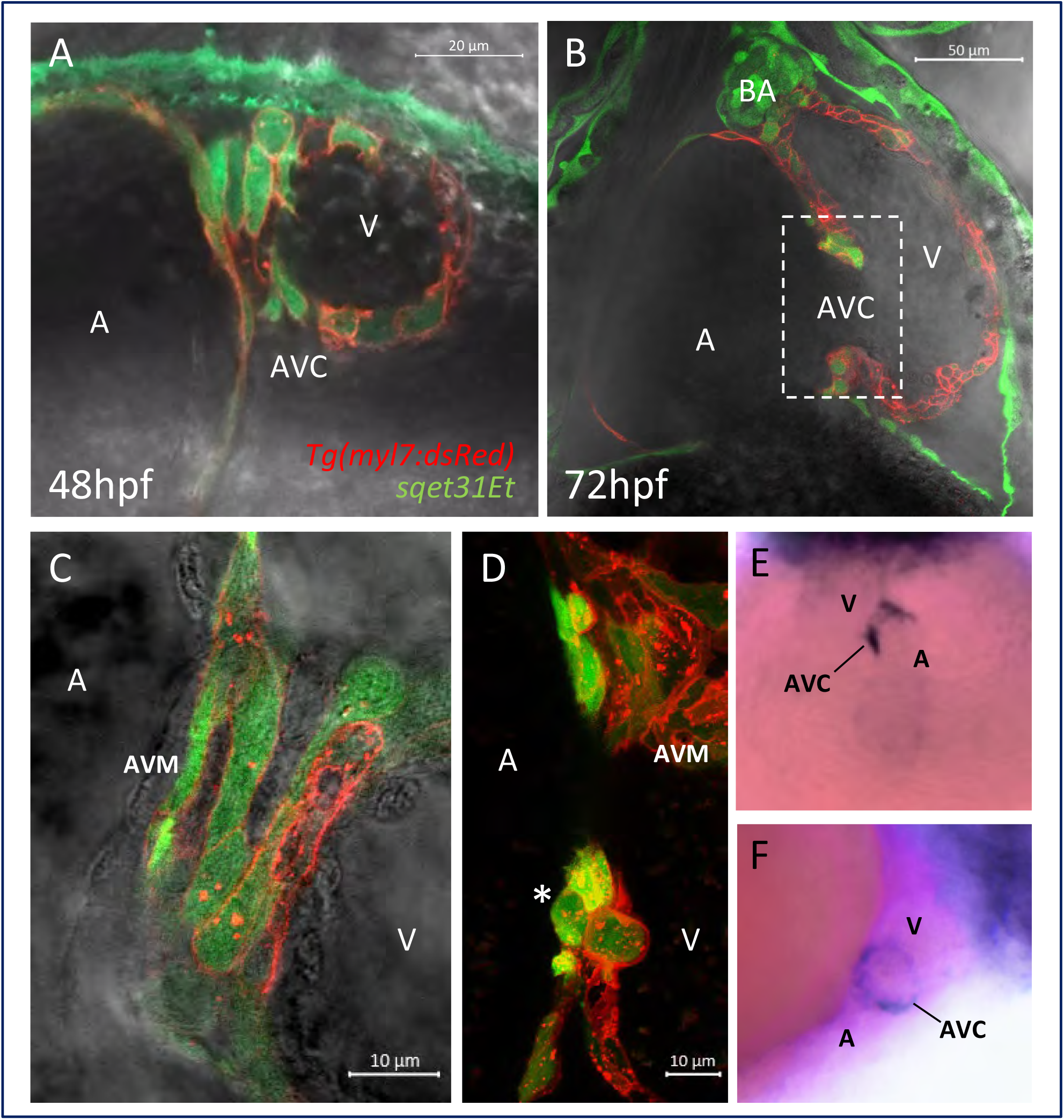
EGFP expression in the transgenic line sqet31Et defines cells of the AVC. (A, B) confocal images showing the AVC region at 48 hpf and 72 hpf. Note the overlap between EGFP and mRFP signals, indicating extensive co-localization and confirming the largely myocardial nature of the EGFP expression domain in *sqet31Et*. (C, D) Close-up of the region marked in panel B at different focal planes showing the surface (C) and lumen (D) of the AVC. Note the group of 3 cuboidal cells protruding into the cardiac lumen at the location corresponding to the developing cardiac cushion (asterisk). (E, F) Whole-mount *in situ* hybridization of *egfp* in *sqet31Et* transgenic embryos at 72 hpf showing enrichment of EGFP expression in the AVC relative to the rest of the heart. E - ventral, D - lateral view, A - atrium, V - ventricle, AVC - atrioventricular canal, BA - bulbus arteriosus, AVM - atrioventricular myocardium.

### Transcriptome profile of the AVC

To characterize the molecular profile of the GFP+ cell population in *sqet31Et,* we isolated these cells using fluorescence-activated cell sorting (FACS) at 48 hpf and 72 hpf (Fig. 2A) and profiled their transcriptome by RNA-seq. The rest of the heart cells, which did not express EGFP, were also collected (GFP-). Average sequencing reads mapping to the *egfp* sequence were considerably higher in GFP+ compared to GFP- samples, confirming the high representation of the EGFP-expressing cell population in the GFP+ samples (S1 Figure B). Principal component analysis (PCA) revealed the tight clustering of replicates and clear separation between sample types (Fig. 2B). Moreover, samples of the same developmental stages clustered closer together, indicating their similarity to each other.

**Figure 2.**
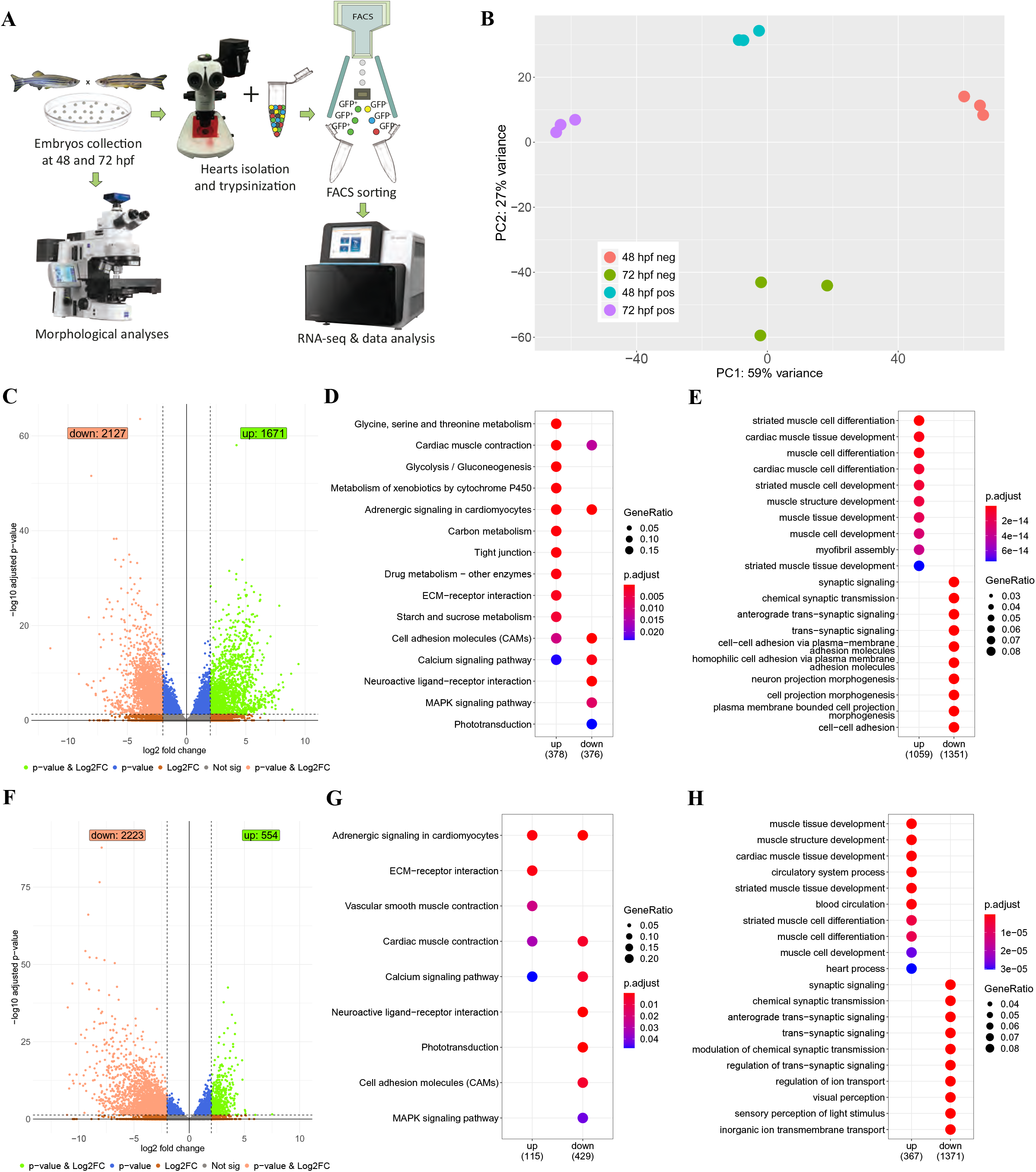
Transcriptome profiling of the GFP+ cells isolated from *sqet31Et.* (A) Scheme of experimental design. (B) Principal component analysis (PCA) on normalized RNA-seq data (regularized log) showing variance between three technical replicates of each sample as well as between samples. (C, F) Volcano plots showing genes differentially expressed between EGFP-positive and -negative cells at 48 hpf (C) and 72 hpf (F). DESeq2 was used to calculate log2FC and padj values. Green spots indicate genes considered as significant (padj < 0.05) with at least two-fold change between groups (log2FC > 2). Accordingly, light red spots represent significant genes with log2FC smaller than -2. (D, G) KEGG pathway enrichment of differentially expressed genes at 48 hpf and 72 hpf. Enrichment analysis was performed on a gene list meeting the following criteria: log2FC >2 or log2FC < -2 and padj < 0.05. Same criteria were used to perform biological process Gene Ontology terms enrichment on up and down-regulated genes (E, H). Top 10 terms for each enrichment analysis were shown.

To identify genes enriched in the GFP+ cell population compared to the rest of the heart, we performed differential expression analysis of the GFP+ against GFP-fractions. In both developmental stages, a total of 3798 and 2777 genes were differentially expressed at 48 hpf and 72 hpf, respectively (absolute log2FC > 2, padj < 0.05), of which 1492 were common for both stages (Fig. 2C, F, S2 Table). GO and KEGG pathway enrichment analyses at 48 hpf revealed that the set of genes overexpressed in GFP+ compared to GFP-cells (enriched genes) was overrepresented for functional terms related to cardiac muscle development and function (“cardiac muscle contraction”, “adrenergic signaling in cardiomyocytes”, “cardiac muscle development”, “cardiac muscle differentiation”, and “calcium signaling pathway”), which supports the myocardial identity of the GFP+ fraction (Fig. 2D, E, S3 Table). On the other hand, functional terms related to cell-cell adhesion (“cell adhesion molecules”, “cell-cell adhesion”) were overrepresented among transcripts overexpressed in the GFP-cell population. At 72 hpf, similar functional terms were overrepresented with the addition of “vascular smooth muscle contraction” term (Fig. 2G, H, S3 Table), which likely corresponds to the initiation of EGFP expression in the BA at this stage.

To assess whether the sorted GFP+ fraction contained AVC cells, we explored the presence of known markers of AVC in our dataset. We established a set of AVC marker genes for zebrafish and mammals by retrieving genes annotated with the term “atrioventricular canal” from the ZFIN (http://zfin.org/) and MGI [54] gene expression databases. Intersection of these known markers with the set of AVC-enriched transcriptome returned 46 and 58 genes in common, which were enriched in GFP+ cells at both 48 hpf and 72 hpf, respectively (Fig. 3A, B; S4 Table). Notably, several of these genes were known to be expressed specifically in zebrafish AVC myocardium, including *bmp4* [36], *wnt2bb* [55], *snai1b* [56], *hey2* [57], and *hnf1ba* [18]. On the other hand, 27 genes which were enriched in the GFP-population overlapped known AVC markers at either or both developmental stages. These included *id4* and *notch1b* reported to be expressed in the endocardium [41, 58], which suggests that the GFP+ cells in *sqet31Et* are less likely to be endocardial. Whole mount *in situ* hybridization of eight selected AVC-enriched transcripts with no previous heart expression data further confirmed their expression in the AVC except for one, *si:dkey-57k2.6*, which is expressed only in the BA. Another transcript, *si:dkey-164f24.2*, was expressed in the whole heart (S2 Figure). Therefore, utilizing the *sqet31Et* transgenic line to specifically enrich for the GFP+ cell population, our RNA-seq analysis revealed the transcriptome mainly representing the AVC, with contribution from the BA.

**Figure 3.**
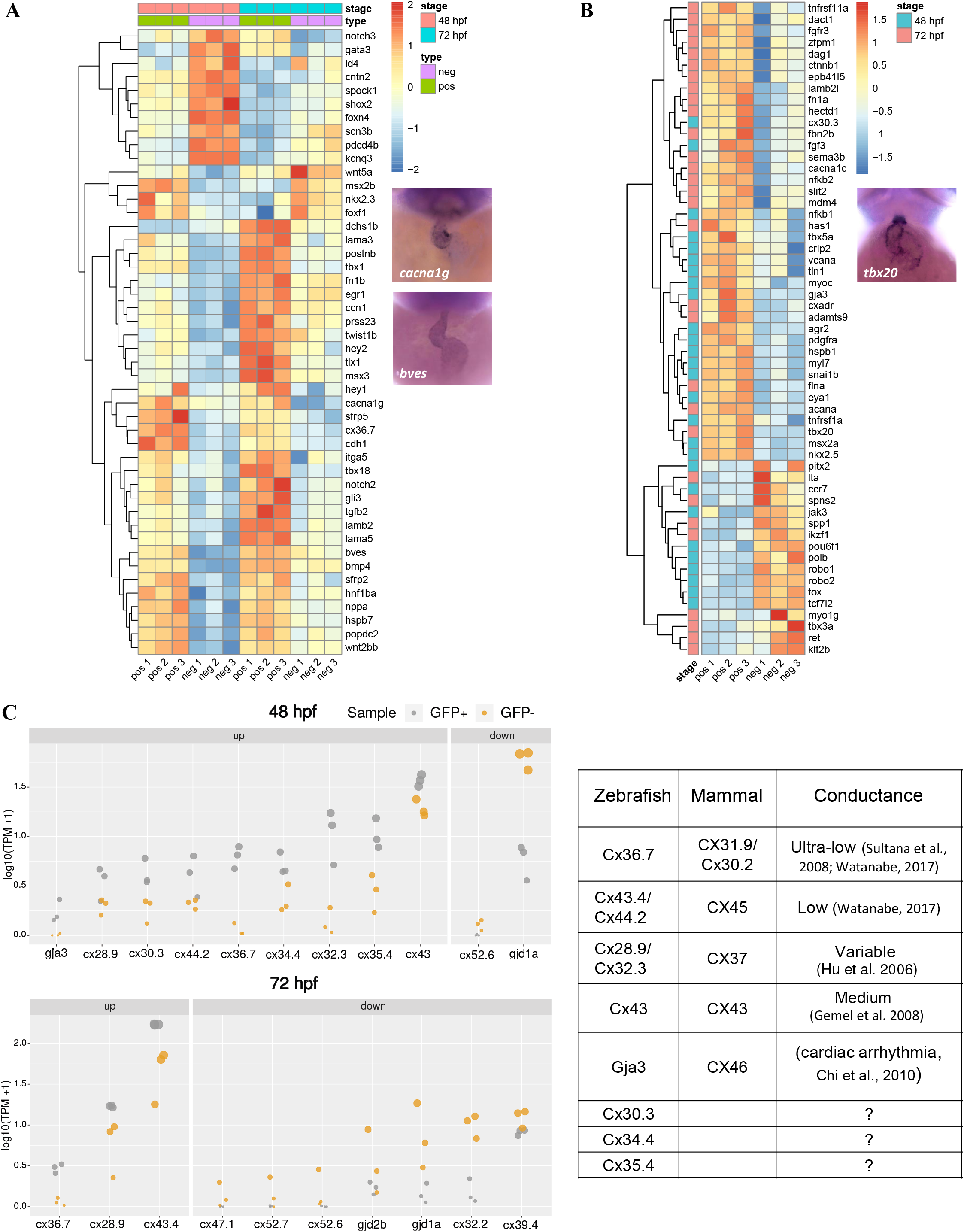
AVC gene signatures are enriched in *sqet31Et* EGFP-expressing cells. Signatures were retrieved from MGI and ZFIN databases and used to identify molecular markers associated with AVC expressed in the studied dataset (padj < 0.05). (A) Heatmap depicts the dynamic of changes of known molecular AVC signatures that are in common across the developmental stage. (B) AVC markers uniquely expressed in either 48 hpf or 72 hpf stage. (C) Expression (in log10(TPM +1)) of genes encoding connexins, components of gap junctions which confer conductance properties between cells, in GFP+ and GFP-cells at 48 hpf and 72 hpf. Mammalian homologs of each connexin gene and their known conductance properties are described in the accompanying table.

### AVC gene expression profile shows signatures of AV pacemaker

Our transcriptome analyses revealed several key features of the zebrafish AVC. Firstly, a hallmark feature of mammalian AVC myocardium is its slow conduction in contrast to chamber myocardium. Among the transcripts enriched in the GFP+ cell population, we found 10 encoding various connexins. These consisted of *cx36.7* (ortholog of human CX31.9 and murine Cx30.2; [59, 60]) and *cx43.4/cx44.2* (ortholog of human CX45; [60]), whose mammalian orthologs form low or ultra-low conductance gap junctions in the AV node [13,14,61]. These were enriched in the GFP+ population at both developmental stages, while other connexin transcripts were enriched only at 48 hpf (padj < 0.05). The latter group consisted of *cx28.9/cx32.3* (ortholog of human CX37; [60]), *cx43*, and *gja3/cx46* (Fig. 3C; S5 Table). Among those with the highest fold-change between the GFP+ and GFP-cell populations were *cx36.7*, *gja3/cx46*, and *cx32.3* at 48 hpf and *cx36.7* at 72 hpf. Loss of function of mammalian CX46 leads to cardiac conduction disorders, while the loss of *gja3/cx46* in the zebrafish mutant *dco* causes defects in heart morphology and ventricular conduction pattern [62]. The enrichment of genes encoding connexins able to form low conductance gap junctions likely reflects the conduction delaying property of the AVC region. Besides those known for their role in cardiac conduction, transcripts encoding other members of the connexin family [Cx30.3 (CX30), Cx34.4 (CX30.3) and Cx35.4 (CX31)] were also enriched in the GFP+ cell population. These have not been previously implicated in heart or pacemaker function and are candidates for further investigation.

Besides delaying electrical conduction between the atrium and ventricle, the mammalian AV node also possesses intrinsic pacemaker activity [3, 6]. We thus searched amongst the AVC-enriched gene list for those known to be expressed in the AV node or associated with pacemaker development and function (S6 Table; [63–66]. Confirming previous reports [28], the expression of *hcn4* was observed in the GFP+ cell population at both 48 hpf and 72 hpf (S3 Figure; S6 Table). In addition, genes encoding zebrafish orthologs of transcription factors known to be involved in mammalian pacemaker cell specification [67–69] were expressed in the GFP+ population (*tbx18, shox2, tbx2a, tbx2b,* and *tbx3*; S3 Figure, S6 Table). In mammalian CCS, the transcription factors Tbx2 and Tbx3 are known to repress the expression of the chamber-specific Cx40 [67, 70]. In agreement with this, *gja5a/b* (the zebrafish ortholog of CX40), was not AVC-enriched. It has been shown that *nkx2.5* is expressed in all myocardium, but slightly higher in the AV conduction system [64]. Similarly, we found that *nkx2.5* was enriched in GFP+ cells compared to GFP-at 48 hpf (S3 Figure, S6 Table). Taken together, the transcriptome of AVC myocardium reveals conserved features to that of the mammalian AV node in terms of expression of genes linked to slow conductivity, automaticity, and molecular mechanism for AV node specification. Therefore, our results support the notion that the AVC is a homologous structure to the mammalian AV node and suggest that the core network leading to the specification of the AV conduction system is conserved between mammals and fish.

### Defect of the primary pacemaker in the SAN reveals spontaneous activity of the AVC pacemaker

The expression of *hcn4* and other markers in AVC, suggesting its homology to the mammalian AV node, led us to question whether the zebrafish AVC possesses inherent pacemaking activity as does its mammalian counterpart. To this end, we utilized the *isl1* K88X mutant (*isl1*^sa29^), which has been shown to exhibit a defective SAR pacemaker function that manifests as sinus pauses and bradycardia [21, 25]. We observed that, apart from losing the expression of *fhf2a* (Fig. 4A, B), *bmp4* (Fig. 4C, D), and the pacemaker marker *hcn4* (Fig. 4E, F) in the sinus venosus, the *isl1* mutant was also devoid of EGFP-positive cells at the SAR, but not the AVC, as shown by analysis of *sqet33mi59BEt* and *sqet31Et*, respectively (Fig. 4G, H). Hence, we confirmed that the reduced number of cardiomyocytes at the venous pole in *isl1^-/-^* observed previously [25] resulted from the absence of the pacemaker cells-containing SAR. It is worth noting that the *isl1*^-/-^ is the only vertebrate mutant that shows a complete lack of pacemaker SAR cells. In support of this, Isl1 overexpression resulted in a small increase in the fluorescence intensity of EGFP-expressing pacemaker cells in *sqet33mi59BEt* at 40 hpf (Fig. 4I, J).

**Figure 4.**
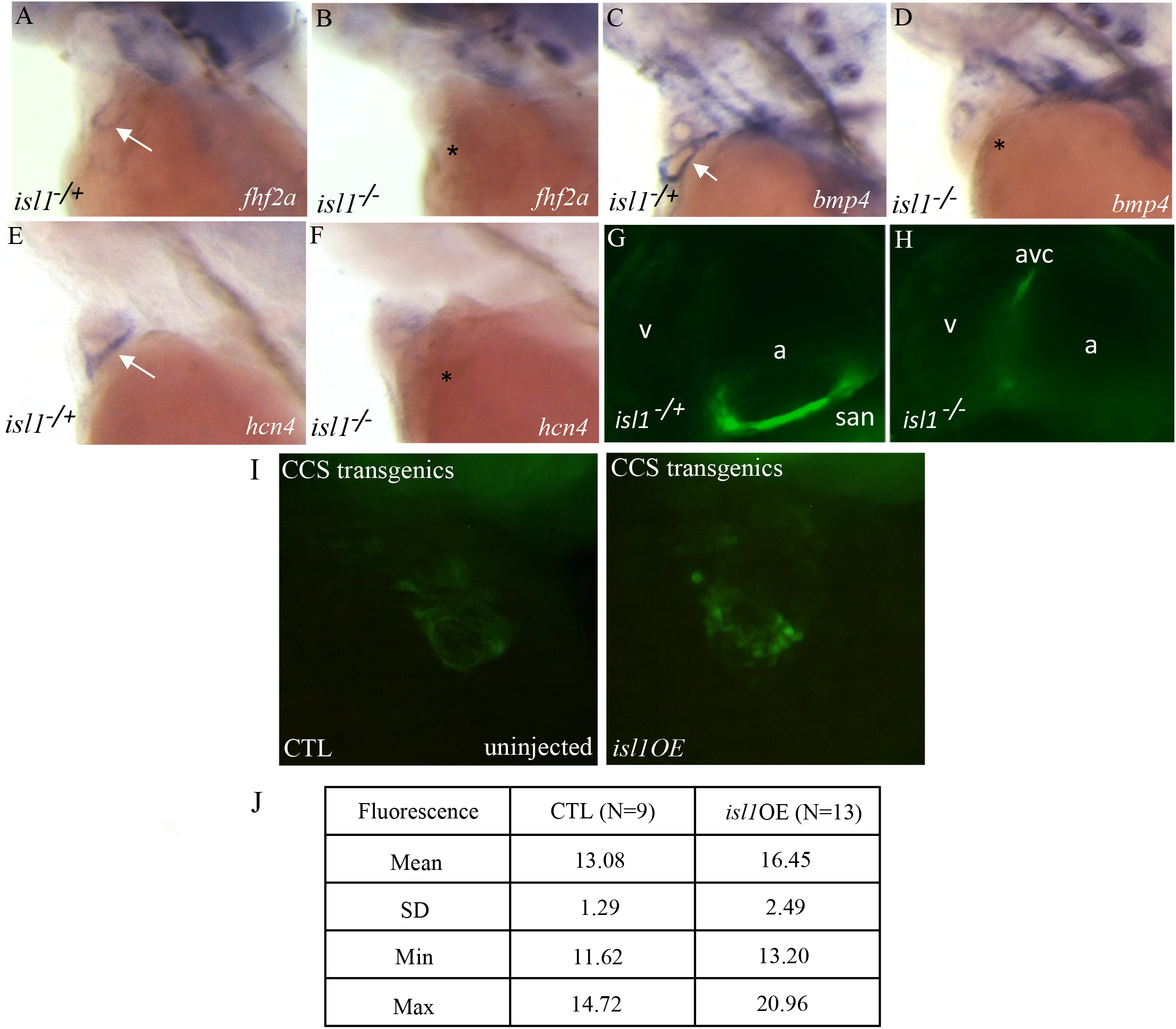
The absence of the pacemaker ring in *isl1* mutant causes the loss of expression of *fhf2, hcn4, bmp4.* (A-H) Expression pattern (labeled with arrow) of *fhf2a* (A, B), *bmp4* (C, D), *hcn4* (E, F) and EGFP (G, H) in *isl1* siblings and mutants. (A-F) - WISH, (G, H) - confocal microscopy of CCS in the sibling and *isl1* mutant. (I) Representative image of CCS transgenics control embryo and *isl1* overexpressed embryos at 40 hpf. (J) The table presents the mean, standard deviation, and minimum and maximum grey value (determined in Image J) as the measured EGFP fluorescence intensity of the pacemaker cells at 40 hpf. The numbers (N) of embryos used for measurement is indicated.

Despite the complete absence of the SAR pacemaker, Isl1-deficient hearts contract, albeit inefficiently and irregularly, with long pauses (Fig. 5A, C, Supplementary movies 1 and 2). This suggests the existence of alternative origins of automaticity that triggers the initiation of cardiac contractions. To investigate whether the AVC could generate electrical impulses independent of the SAR, the Isl1 antisense morpholino [71] was injected into 1-4 cell stage zebrafish embryos expressing the genetically encoded voltage-sensitive fluorescent protein Mermaid, *Tg(myl7:mermaid)*. This allowed direct observation of cardiac electrical conduction patterns. Similar to *isl1* mutants, about 66% of Isl1 morphants showed sinus pauses (6 of 9 morphants *vs* 4 of 6 mutants), and all showed bradycardia. The heart rate of Isl1 morphants was 80.2±15 beats per minute (bpm) at 48 hpf (mean ± SEM., n=6) and 141,2±10.3 bpm at 72 hpf, (n=6), which is significantly lower than that in controls (176±5.7 bpm at 48 hpf, (n=7) and 229±6.6 bpm at 72 hpf, (n=18)) (Fig. 5A). Increased variability in heartbeat duration was noted in the Isl1 morphants as well.

**Figure 5.**
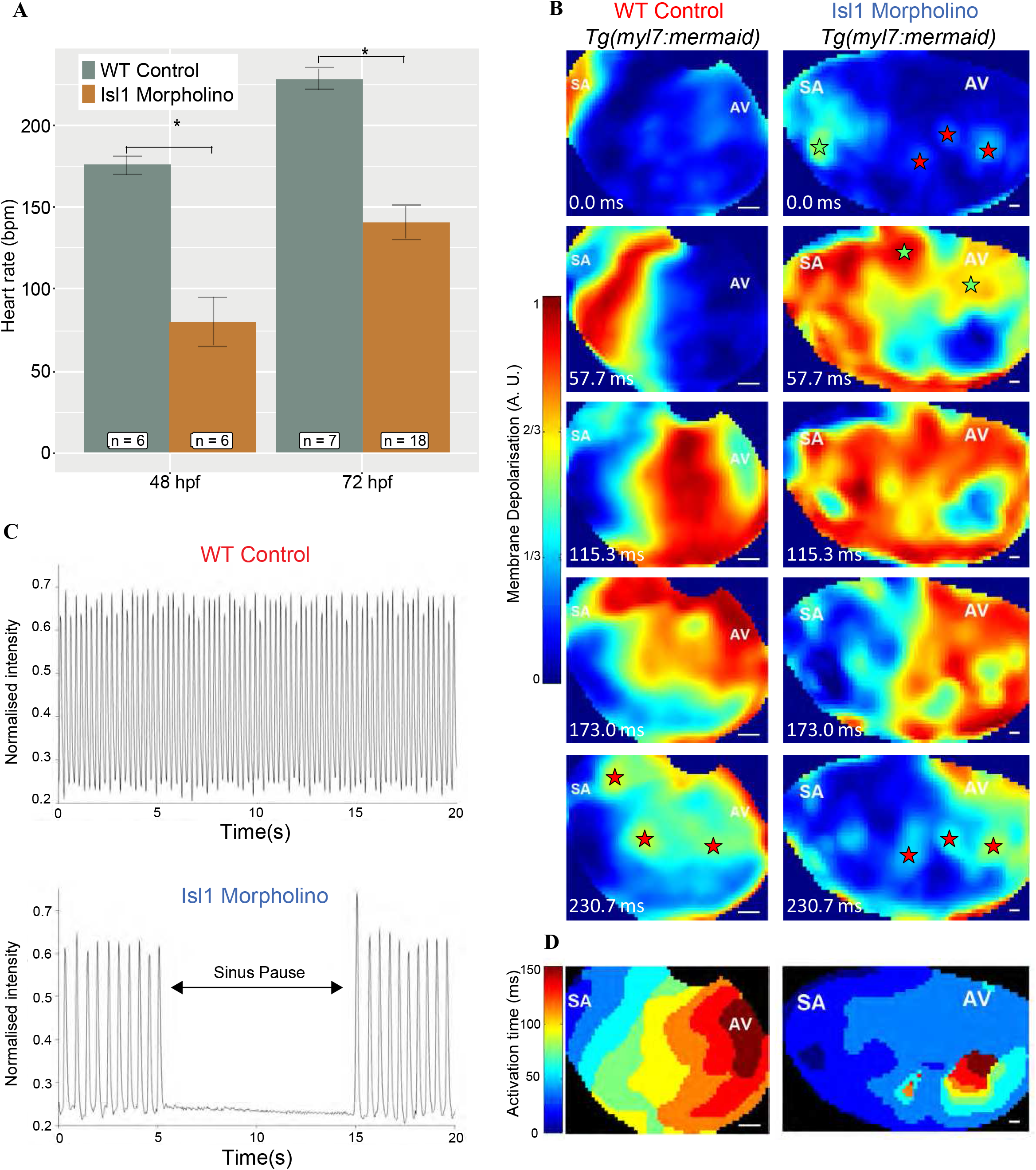
Effects of morpholino knockdown of isl1 on electrical activity of the atria in zebrafish larvae. (A) Heart rate at 48 hpf (left) and 72 hpf (right) in WT control (red) and *isl1* morphant (blue) zebrafish, showing slowed heart rate in *isl1* morphants. \**p*<0.001. (C) Videographic analysis of the heartbeat in WT control (top) and *isl1* morphant (bottom) 48 hpf zebrafish, showing slowed heart rate and sinus pauses (during the period indicated by the arrow) in the *isl1* morphant. (B) Sequence of video frames showing electrical activation of the atria in WT control (left) and *isl1* morphant (right) 48 hpf *Tg*(*myl7:mermaid*) zebrafish, showing normal activation (from the sinoatrial node region [SA] to the atrioventricular [AV] junction) and sites of latest activation/repolarisation (indicated by red stars) in the WT control and abnormal activation in the *isl1* morphant (sites of early/ectopic activation in the morphant zebrafish indicated by green stars). (D) Isochronal activation map of the atria in WT control (left) and *isl1* morphant (right) 48 hpf zebrafish derived from the video represented in B.

Optical mapping showed that in control hearts, the excitation wave front travelled uniformly across the atrium from the SAR towards the AVC (Supplementary movie 1; Fig. 5B). In contrast, in Isl1 morphants, atrial excitation, although predominantly originating from the SAR region, became uncoordinated (Supplementary movie 2; Fig. 5B). During the sinus pause, several stationary centers of automaticity distributed from posterior to anterior were detected, with the SAN-associated location showing the highest activity. Interestingly, the location associated with AVC became more active near the end of the atrial contraction (Supplementary movie 2, Fig. 5B, asterisk), presumably when triggering the consecutive phase of the contraction cycle in the ventricle. The activity of the AVC location became even more pronounced during the periods of sinus pause. The loss of coordinated atrial excitation and conduction in Isl1 deficient embryos therefore suggests that the SAR is the primary pacemaker required for coordination of atrial excitation. However, coordinated ventricular excitation can be induced by the electrical activity of the AVC, in particular when no wave of excitation from the SAR drives the heartbeat. Inherent AVC pacemaker rate is lower than that of the SAR.

### Comparison between the AVC and SAR transcriptomes reflect distinct electrophysiological properties

The observation that both SAR and AVC regions of the zebrafish heart possess pacemaker activity, albeit at different inherent rates, led us to question the molecular nature underlying their distinct properties. To this end, we compared the transcriptome of the AVC with that of the SAR [52] to identify differentially enriched genes. We re-analyzed the SAR dataset to identify SAR-enriched genes and intersected this with our AVC-enriched gene list (S2 Table), obtaining a total of 1516 AVC-unique and 701 SAR-unique genes, and 450 genes common between the two (S7 Table). Notably, several genes encoding proteins involved in ion transport, cell junction formation, and extracellular matrix were differentially enriched between the AVC and SAR.

Interestingly, the expression of *hcn4* was significantly higher in the SAR, while not significantly higher in the AVC, compared to the rest of the heart. This may reflect the role of SAR as the dominant pacemaker (S7 Table). In addition to *hcn4*, several other transcripts encoding various ion channels were enriched in both the SAR and AVC, notably, the T-type calcium channel Cacna1g, which is necessary for mammalian pacemaker activity in both SA and AV nodes [72]. Those genes enriched in AVC and not the SAR include *trpm4* encoding a Ca^2+^-activated nonselective cation channel, which mediates cell membrane depolarization [73]. TRPM4 is implicated in human progressive familial heart block type I characterized by cardiac conduction blockage downstream of the AV node [74]. Another notable example is *cacna1c*, whose human ortholog is associated with the Wolff-Parkinson-White syndrome [75], a condition affecting the AV conduction system [76]. Other genes, including *kcnq1.1*, *kcne4*, and *atp1b1a*, possess human orthologs associated with the maintenance of QT interval [77–79].

Despite having some common properties, the SA and AV nodes perform distinct functions. The SA node serves a primarily pacemaking function, while the AV node is mainly specialized to delay electrical propagation between the atrial and ventricular chambers [11]. Therefore, while both regions express partially overlapping, mostly low-conducting, gap junction proteins [11, 80], the AV node is particularly enriched in Cx30.2 and Cx45 [61,81,82]. In line with this, *cx36.7,* the zebrafish paralog of *Cx30.2*, was enriched in the AVC but not the SAR (S7 Table). On the other hand, *cx43.4*, a paralog of *Cx45,* was enriched in both the SAR and AVC (S7 Table). Low electrical coupling is also a necessary property within the definitive pacemaker cells of the SA node to prevent inhibitory interference from the surrounding working myocardium, which is more hyperpolarized [11]. Collectively, the overall differences in ion channel, cell adhesion, and extracellular matrix composition enriched in the SAR and AVC likely underlie their distinct electrophysiological properties.

### Developmental signaling pathways dynamics suggests signaling interaction between AV myocardium and endocardial cells during valve formation

Besides hosting the AV node, the AVC is also the site of valve formation. We identified transcripts encoding genes involved in the EMT process enriched in the GFP+ compared to GFP-cell population (Fig. 6A, B; S8 Table). These include 22 genes enriched at both developmental stages and 28 genes enriched only at either the 48 hpf or 72 hpf stage. Members of the TGF-β signaling pathway were enriched in GFP+ cells at both developmental stages (*tgfb2*, *tgfb1a, smad1, smad6b, and smad9*) (Fig. 6C - E). In addition, the GFP+ enrichment of *tgfb3, tgfbr2b,* and *smad6a* in GFP+ cells was specific to the 48 hpf stage (Fig. 6C). This is in line with the observation in mammalian endocardial cushion formation, where various TGF-β ligands are expressed in different cell populations of the AVC: TGF-β1 in endocardium, TGF-β2 in myocardial and endocardial cells flanking the cushions, and TGF-β3 in the cushion mesenchyme [83].

**Figure 6.**
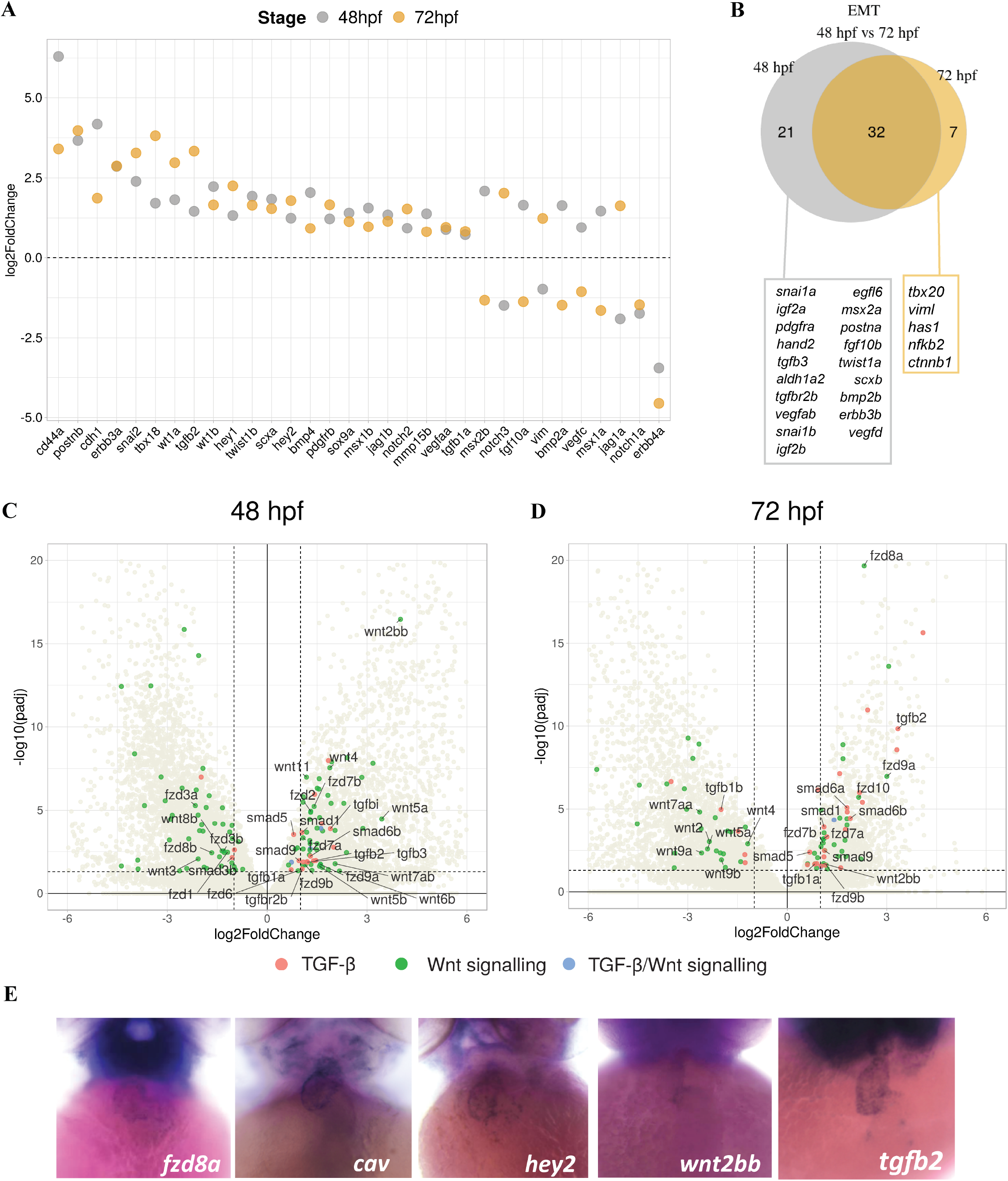
The transcripts of genes involved in EMT and valve development are enriched in the AVC. (A) AVC-enrichment (expressed in log2 fold change between GFP+ and GFP-cells, padj < 0.05) of genes known to regulate EMT at both 48 hpf and 72 hpf stages. (B) Overlap of EMT-regulating genes enriched in GFP+ cells at both stages. (C-D) Volcano plot showing enrichment of components of the TGF-β and Wnt signaling pathways in AVC at both 48 hpf and 72 hpf stages. (E) Whole mount *in situ* hybridization of several AVC-enriched genes of the TGF-β and Wnt pathway components.

Notch signaling activity in the AVC endocardium is necessary for the initiation of the EMT process by activating SNAI1 and SNAI2 expression [84, 85]. Transcripts encoding key components of the Notch pathway were enriched in GFP+ cells at both developmental stages. These include *jag1b*, *hey1*, and *hey2,* and *prss23*, implicated in AV valve formation [86].On the other hand, *jag1a*, *jag2a*, *her6*, *jagn1b*, *notch2*, and *notch3* were enriched at either stage (S2 Table). In contrast, valve endocardial markers *notch1b* and its ligand *dll4* [34–36] were not enriched in GFP+ cells, which further supports the non-endocardial identity of this population. Several components of the canonical Wnt signaling have been reported to be expressed in different cell layers of the mammalian AVC, including *Wnt2*, *Fzd2*, and *Lef1* in the cushion mesenchyme and *Wnt4* and *Wnt9b* in endocardial endothelium [87]. In agreement with this, we found that, in the zebrafish AVC, *wnt2bb* was enriched in GFP+ cells at both stages, while *wnt7ab*, *wnt6b*, *wnt5a/b*, and *wnt4* were enriched only in 48 hpf (Fig. 6C - E). On the other hand, *fzd7a/b* and *fzd9a/b* encoding Wnt signaling receptors were enriched in both stages, *fzd2* and *fzd6* enriched only at 48 hpf, while *fzd6*, *fzd8a,* and *fzd10* only at 72 hpf (Fig. 6C-E). Whereas *dkk1a/b* and *dkk2* encoding Wnt antagonists were enriched only at 48 hpf. Interestingly, we observed that at 72 hpf, except for *wnt2bb*, no transcripts of Wnt ligands were enriched in GFP+ cells, while transcripts encoding several Frizzled family receptors and downstream components were enriched (Fig. 6D). To visualize Wnt signaling activity in the AVC region, we crossed the *sqet31Et* transgenic line with the Wnt reporter line *Tg(7x TCF-Xla.Sia:NLS-mCherry)* [88]. In agreement with previous reports of the presence of Wnt signaling in the AVC myocardium [40, 89], we observed that both the non-EGFP-expressing endocardium and EGFP-expressing cells had Wnt signaling activity at all developmental stages (S4 Figure).

The process of valve development is reflected in the dynamics of signaling pathway activity occurring between the early and late stages of AVC development. We found a total of 5877 genes differentially expressed at 72 hpf compared to 48 hpf (padj < 0.05; -1 < log2FC > 1; S5 Figure, S9 Table). Notably, GO terms related to TGF-β, Wnt, and Notch signaling pathways were overrepresented at 72 hpf (S10 Table). Many members of these three signaling pathways exhibited dynamic expression between 48 hpf and 72 hpf (S5 Figure; S11 Table). Collectively, our observations uncover the dynamic expression of various components of the TGF-β, canonical Wnt, and Notch signaling pathways in the AVC myocardium, which likely reflects their role in the ongoing AVC patterning and valve development.

### AVC-enriched genes are associated with human congenital heart defects related to CCS, valves and septa

We identified the human orthologues of AVC-enriched genes and interrogated them for any association with clinical phenotypes related to ClinVar terms: “arrhythmia”, “AV block”, “long QT syndrome”, and “conduction”. Our analysis revealed a total of 91 and 60 unique genes associated with these four ClinVar terms at 48 hpf and 72 hpf stages, respectively (S12 Table). Specifically, the disease conditions represented by these terms included more general forms of cardiac arrhythmia such as atrial fibrillation, sick sinus syndrome, abnormal QT interval, and Brugada syndrome, as well as those conditions specifically associated with defects of the AV conduction system or downstream effects such as heart block [90], Wolff-Parkinson-White pattern [76], and supraventricular tachycardia [91]. The latter group included *trpm4* and *cacna1c,* which were enriched in both SAR and AVC, as well as *mybpc3, smyhc2, hrc, dspa, myh7l, zgc:86709, lmna, snta1*, and *ttn.2*.

As the AVC is also the site where the endocardial cushion and valve develops, we expected to find associations between AVC-enriched genes and human valve and septal defects. We searched the human orthologues of AVC-enriched for overlap with ClinVar terms containing “tricuspid valve”, “AV valve”, “mitral valve”, and “valve in general”. In total, 115 and 93 unique genes were associated with these terms at 48 hpf and 72 hpf, respectively (S13 Table). In addition, we also searched for those associated with the ClinVar term “septal defect” and obtained 66 and 55 unique genes associated with the term at each stage, respectively (Supplementary Table 14). In the adult human heart, the AV node is embedded into the interatrial septum [3]. Given that the endocardial cushions are involved in the formation of the AV valves and septa, it comes as no surprise that the defects of interatrial septum could be linked to defects in cardiac conduction. In fact, a number of genes were commonly associated with ClinVar terms “cardiac conduction” and “valve” (S14 Table). For example, *tbx5a*, whose human orthologue TBX5 causes the Holt-Oram syndrome characterized by congenital heart malformation due to variable atrial and ventricular septal defects as well as heart conduction defects [92, 93]. Another notable example is *smyhc2*, whose human orthologue MHY6 is associated with both atrial septal defect and sick sinus syndrome [94, 95]. Other examples include *cacna1c*, *ttn.2*, *snta1*, *lmna*, *dspa*, and *mybpc3.* Taken together, the overlap of a large number of AVC-enriched genes with human heart conditions related to CCS and valve/septal defects suggests our transcriptomics data as a valuable resource for studying these diseases.

### The bulbus arteriosus transcriptome contributes to a set of differentially expressed genes at 72 hpf

Among the GO terms associated with a set of genes enriched at 72 hpf is “Vascular smooth muscle contraction”. This term likely reflects the expression of EGFP in the BA at 72 hpf. The BA is a teleost-specific structure, which exists in place of the outflow tract of higher vertebrates [53]. Unlike the outflow tract, which is rich in cardiomyocytes, the BA is mainly composed of smooth muscle [96]. In the *sqet31Et* transgenic line, the BA-associated GFP expression was observed at 72 hpf (Fig. 7A). Accordingly, we observed the enrichment of transcripts encoding several smooth muscle markers, including smooth muscle light chain kinase *mylka*, in the 72 hpf dataset (Fig. 7B). The BA dampens blood pressure fluctuations occurring during different phases of the cardiac cycle [97]. The gene encoding Elnb, which is known to promote the differentiation of smooth muscle cells [53], is expressed in the BA starting from 72 hpf (Fig. 7B). In agreement with this, we found that *elnb* and its paralog *elna* were among genes enriched at 72 hpf compared to 48 hpf. Moreover, we also found the enrichment of other genes previously implicated in maturation and function of elastin, including *ltbp3* and *fbln5* (Fig.7B) [53,98,99]. To date, only one study reported the transcriptome profile of the BA in the adult zebrafish [100]. We found that 56 out of the 59 BA-expressed genes identified in their study were enriched in GFP+ cells at 72 hpf (S15 Table). Taken together, our results suggest that the 72 hpf transcriptome of GFP+ cells defines genes expressed in the AVC and BA. Nevertheless, the distinctive tissue composition of the BA compared to AVC myocardium allows for segregation based on this criterion. The expression of EGFP in the AVC and BA of *sqet31Et* transgenic line adds to a list of common markers of these cell lineages.

**Figure 7.**
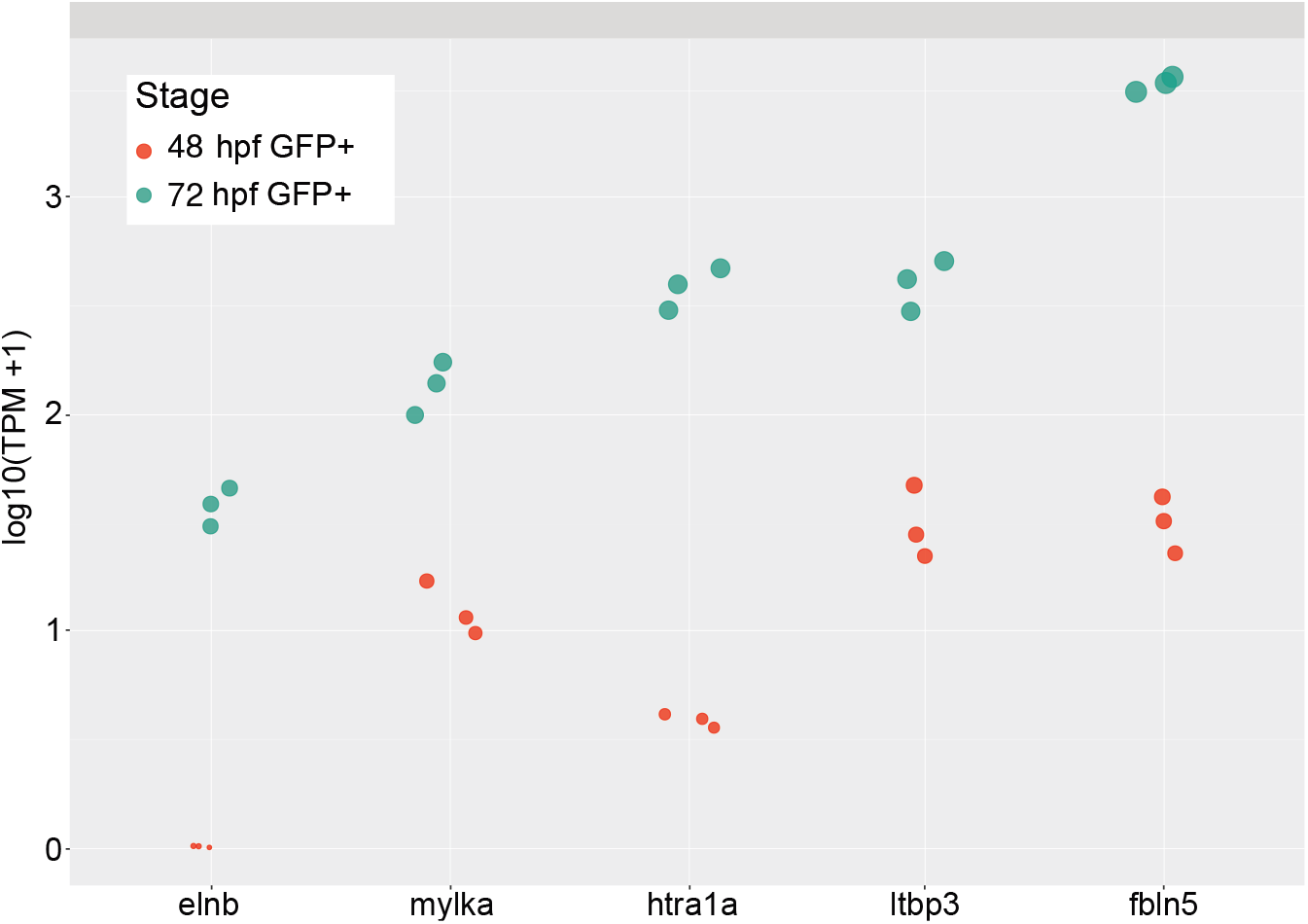
Bulbus arteriosus constitute the 72 hpf transcriptome. (A) Confocal image of the heart in double transgenic line *sqet31Et* x *Tg(myl7*:*mRFP*) at 72 hpf showing additional EGFP expression domain in the bulbus arteriosus (BA). (B) Genes previously known to be expressed in the BA are enriched at 72 hpf compared to 48 hpf. In the zebrafish, the BA is composed of smooth muscle and is rich in elastin and contractile proteins. Note the high enrichment of *elnb* at 72 hpf as compared to 48 hpf.

## Discussion

The AVC plays two major roles. First, it is part of the CCS which serves as the site where the propagation of electrical impulses is delayed, allowing consecutive contraction of the atrium and ventricle. In mammals, this function is performed by the AV node. Second, the AVC is also the site where the heart valve develops, involving various signaling pathways and cellular rearrangements occurring between different tissue layers. The study of the AVC has been challenging due to the lack of specific molecular markers defining this region. Available data relied on methods based on histological sections [101], which lacks the ability to isolate specific cell types. Nevertheless, it is known that distinct structures of the AVC, such as the pacemaker cells and cardiac valve tissue, express unique combinations of marker genes which can be used to distinguish them.

The transgenic line *sqet31Et* provides the necessary level of specificity, which allows the enrichment of AVC myocardial cells by FACS. By this strategy, in combination with transcriptome analyses, we demonstrate that the zebrafish AVC myocardium possesses hallmarks of the mammalian AV node. The AVC transcriptome is characterized by high expression of mRNA encoding low conductance connexins *cx36.7* and *cx43.4*, as well as the T-type calcium channel *cacna1g* and pacemaker hyperpolarization-activated channel *hcn4*. All these factors are known to define the AV node and pacemaker activity in the mammalian heart [61,72,81,82,102]. The conserved features also extend to the expression of transcripts encoding the core pacemaker transcriptional network consisting of Tbx2a/2b/3/18 and Shox2 transcription factors.

Previously, the existence of the CCS in zebrafish has been supported by optogenetic studies [27, 29] with other evidence suggesting that the endocardium and hemodynamic stimulation play an important role in its development [51,58,103]. The primary SA pacemaker has been relatively better characterized, and it has been shown that its activity depends on Isl1 [21]. Using the *sqet33mi59BEt* transgenic line, we show that the loss of Isl1 in mutants abolishes the SAR, which harbors the primary pacemaker activity. Furthermore, analysis of electrical conduction patterns in *isl1* morphants revealed that, despite being disorganized, atrial activation wave generally still progresses from the SA to AV. This suggests that Isl1 is not the determinant of primary pacemaker function but rather acts in coordinating atrial activation. However, we cannot rule out the possibility that the *isl1* knockdown did not abolish Isl1 function completely. In either case, the lack of an organized atrial activation pattern affects overall cardiac contraction, indicating that the coordinated signaling from the SAR and its propagation play a crucial role in coordinating heart contraction.

Interestingly, during the pause in heart rhythm in Isl1-deficient embryos, the observed increase in AVC activation demonstrated that it possesses inherent automaticity, enabling it to independently excite when a weakened signal from the primary SAR pacemaker is not sufficient to drive heart contractions. This corroborates previous observations in adult zebrafish heart of spontaneous electrical activity at the AV region following surgical uncoupling of the ventricle from the atrium [27]. Comparison of the transcriptome profiles of the SAR and AVC, while revealing common markers of pacemaker activity such as *cacna1g* and *cx43.4*, also indicated differences that would affect electrical properties, such as the enrichment of distinct types of ion channel, gap junction, and extracellular matrix components. This further supports the notion that, despite their shared ability to act as a pacemaker, the SAR and AVC performs different inherent functions.

The mammalian AV node consists not only of definitive pacemaker cells, but also fibroblasts, macrophages, and ECM, which provide electrical insulation around the electrically conducting AV node, which forms the only point of continuity between cardiac chambers [2, 42]. Electrical impulses from the AV node are further propagated by means of an intricate network of His/Purkinje fibers, which link to the thick myocardial tissue throughout the whole ventricle [43]. In contrast, the 2-chambered heart in most fishes resembles the mammalian embryonic heart tube, where electrical current is propagated from one end to the other by means of electrical coupling of cardiomyocytes without a specialized CCS [2]. In fact, the hearts of ectothermic animals do not possess any insulating fibrous structure, although the slow conducting muscles of the AVC is present [20]. Moreover, teleost hearts are not known to possess any defined Purkinje fiber network, and conduction function is served by the ventricular trabeculae, which form myocardial continuity between AVC and apex of the ventricle [104]. Therefore, it is reasonable to assume that conduction function could be served by an equally simplified structure, in which a subset of cardiomyocytes performs pacemaking function and at the same time express additional attributes, which enable it to slow down electrical propagation. Intriguingly, in an earlier analysis of the CCS transgenic lines [51], it was observed in the embryonic heart that cells of the SAR send processes into the AVC, which appeared as a network connecting the two structures. It is therefore tempting to speculate that perhaps a previously uncharacterized structure or cell type may exist in the zebrafish, which facilitates fast conduction between the SAR and AVC.

Besides serving as part of the CCS, the AVC is also the site where the heart valves and septa develop. This process involves several signaling pathways and cellular rearrangements occurring in different tissue layers. We found that the AVC was enriched for transcripts encoding regulators of EMT, which is a hallmark of endocardial cushion and valve development. Notable examples include *postnb*, which is one of the most highly enriched EMT-related genes at both developmental stages. Its paralog, *postna,* was also enriched in 48 hpf. In mammals, Postn is known to be expressed in the developing endocardial cushion, where an active EMT process is ongoing, but not in cardiomyocytes [105]. Postn acts as an adhesion molecule marking mesenchymal cells in the developing AV valve leaflets [106]. The higher expression of *postna* and *postnb* in GFP+ cells therefore suggests the presence of mesenchymal cells of the valve within this population and suggest an overall heterogeneity of AVC cell populations marked by EGFP expression in the *sqet31Et* transgenic line. Furthermore, components of major signaling pathways, including Wnt, Notch, and TGF-β, which were implicated in endocardial cushion and valve development [33, 34], were differentially expressed at 48 to 72 hpf. Canonical Wnt signaling is known to play multiple roles in valve development, including regulation of AVC maturation and establishment of its electrical properties upstream of Tbx3 [89]. We observed that Wnt signaling activity is present not only in AVC endocardial cells, but also in the myocardium. This is in line with the observation in mammalian heart that Wnt ligands originating from the endocardium act as a paracrine signal towards the myocardium, which in turn induces endocardial EMT [34]. In the adult zebrafish heart, a compact group of HCN4-positive cells is embedded within the musculature of the AV valves [27]. The close association between the valve tissues and pacemaker cells is reflected in our transcriptome and adds to the heterogeneity of cell types present within this region. Currently, the bulk RNA-seq approach utilized here does not allow us to distinguish between the various cell populations, nor to clearly demarcate the concurrent developmental processes within the AVC region. Further studies at the single-cell level in both embryonic and adult zebrafish hearts are needed to reveal the true cellular diversity of this structure and more accurately characterize the organization of the CCS in zebrafish heart.

Besides the AVC, the zebrafish *sqet31Et* transgenic line expresses EGFP in the bulbus arteriosus. This expression pattern is driven by the activity of a yet unknown enhancer [50]. The co-expression of EGFP in the BA and cell populations within the AVC region suggests a unifying regulatory principle governing the specification of different cell types present at different spatiotemporal contexts. Some regulatory TFs (Tbx3) are expressed in the AV node and the mesenchyme of the outflow tract, suggesting a similarity of the developmental mechanism in these cell lineages [107]. This poses an interesting question on gene regulation by the regulatory element(s) driving this expression pattern in the sqet31Et transgenic line. For now, it is challenging to identify this enhancer due to the insertion of the enhancer trap construct in genomic repeat regions. However, with the increasing availability of long read sequencing methods, it may be possible in the near future to map the insertion site and trace the identity of this enhancer.

Collectively, our results establish that the zebrafish AVC possesses molecular and physiological hallmarks of a secondary pacemaker, similar to that of the mammalian AV node, in terms of automaticity, low conductance properties, and conserved expression of developmental genes. The partially overlapping expression profiles of genes encoding ion channels and connexins likely underlies the distinct conduction functions between the SAR and AVC. In addition, the dynamic expression of signaling pathways implicated in the ongoing valve development illustrates the role of the AVC in both electrophysiological as well as structural separation between the heart chambers. The AVC transcriptome data generated in this study will enrich our knowledge of molecular factors, as well as identify potential new candidates, implicated in cardiac conduction and valve development.

## Methods

### Zebrafish

Wild-type, *sqet31Et* and *sqet33mi59BEt* enhancer trap [50, 51], and other zebrafish lines used in this study: *Tg(myl7:mRFP)* [108], Wnt reporter line *Tg(7xTCF-Xla.Siam:nlsmCherry)* [88], were maintained in the zebrafish facility of the International Institute of Molecular and Cell Biology in Warsaw (license no. PL14656251) in line with standard procedures and ethical guidelines. *Tg(myl7:mermaid)* was generated by injection of a *Tol2-myl7-mermaid* construct (kind gift of Yasushi Okamura), together with transposase RNA, into 1-2 cell stage AB zebrafish embryos followed by screening for fluorescence progeny. The *isl1* K88X mutant (*isl1*^sa29^), and *Tg(myl7:mermaid)* was bred and maintained at the Harefield Heart Science Centre according to the Animals (Scientific Procedures) Act 1986. Embryos were raised in egg water at 28°C, screened for a fluorescence signal in the heart and staged at 48 hpf and 72 hpf based on established morphological criteria[109].

### Heart extraction and fluorescence-activated cell sorting (FACS)

To isolate the heart, embryos were anesthetized with Tricaine (0.16 mg/ml in egg water) and large-scale extraction was performed according to a previously published protocol, with minor adjustments [110]. GFP-expressing hearts were manually separated from remaining tissue under a fluorescence stereomicroscope and collected into 0,5 ml of EDM (L-15/10% FBS). Pools of 300-500 hearts were dissociated with Trypsin-EDTA solution (0.05%) as previously described [111]. A FACS Aria II cytometer (BD Biosciences, USA) was used to enrich GFP positive (fluorescent) and GFP negative (nonfluorescent) heart fractions. Gates for cell sorting were calibrated against dissociated hearts extracted from wild type zebrafish embryos at the respective developmental stages (48 hpf or 72 hpf). On average, FACS yielded 15-25% of GFP+ events of total singlet events (Supplementary Figure 1A).

### RNA extraction

To obtain high-quality total RNA, cells were sorted directly to 500 µl TRIzol™ LS Reagent (Thermo Fisher Scientific, USA) followed by RNA purification and DNase I treatment by means of a Direct-zol™ kit (Zymo Research, USA) according to the manufacturer’s protocol. The Tapestation 2200 and High Sensitivity RNA ScreenTape assay (Agilent Technologies, USA) together with Quantus™ Fluorometer (Promega, USA) were used to assess quantity and quality of total RNA. The average RNA Integrity Number equivalent (RIN^e^) for samples used for downstream analysis was 8.7.

### Library preparation and sequencing

To obtain sequencing libraries, a two-step approach was applied. First, cDNA carrying full-length transcript information was synthesized with SMART-Seq® v4 Ultra® Low Input RNA Kit for Sequencing (TaKaRa Bio, Japan) followed by Nextera XT DNA Library Preparation Kit (Illumina, USA) according to the manufacturer’s guidelines. As previously, Tapestation 2200 and dedicated High Sensitivity D5000 ScreenTape and High Sensitivity D1000 ScreenTape assays were used to validate final cDNA and sequencing libraries, respectively. Final libraries were quantified with KAPA Library Quantification Kit Illumina® Platforms (Kapa Biosystems, USA), followed by paired-end sequencing (2 x 75 bp) performed with Nextseq 500 (Illumina, USA). Libraries were sequenced in triplicate, where a single replicate consisted of GFP-positive and GFP-negative fractions for both developmental stages (48 hpf and 72 hpf), at an average depth of 47 million reads.

### Analysis of sequencing data

FastQC tool v. 0.11.8 [112] was used to assess the quality of obtained raw RNA-seq reads. Minor adapters contaminations were removed by Cutadapt v. 1.17 [113] and RNA-seq reads were mapped to the zebrafish reference genome (GRCz11) using Salmon tool v. 0.9.1 [37] resulting in an average of 75% mappability rate (S1 Figure C). Sequencing reads were further analyzed in R programming language v. 3.5.2 [114], whereas differentially expressed genes were identified by the DESeq2 package [115]. Principal component analysis was performed on normalized reads counts transformed to the log2 scale by *plotPCA* function from the same package. ClusterProfiler v. 3.17.3 [116] was used to calculate the enrichment of both biological processes of Gene Ontology terms as well as KEGG pathways. The *enrichGO* and *enrichKEGG* functions were used with default *pvalueCutoff* and *qvalueCutoff* parameters. The ggplot2 package [117] was utilized for plots generation. All sequencing data have been deposited in the GEO database under accession number GSE160107.

### Confocal imaging

Embryos used for imaging were grown in egg water supplemented with 0.003% 1-phenyl 2-thiourea (PTU) at 24 hpf to prevent the formation of melanophores and pigmentation. Prior to imaging, embryos were anesthetized with 0.02% tricaine (MS-222; Sigma-Aldrich A5040), embedded in 1 % low-melt agarose (Sigma, USA) in egg water, and mounted in a glass-bottom dish before imaging on an inverted confocal microscope (LSM800, Zeiss). Images were further processed with Imaris 8 software (Bitplane).

### Optical mapping of atrial excitation

To visualize excitation in the embryonic heart, a transgenic zebrafish line expressing the FRET-based voltage-sensitive fluorescent protein Mermaid [118] specifically in myocardial cells *Tg(myl7:mermaid)* was used and optical mapping was performed as described previously, with minor adjustments [119]. Injection of morpholino against *isl1* (5′-TTAATCTGCGTTACCTGATGTAGTC-3′) was performed as previously described [71].Embryos were embedded in 1% low melting agarose on a 35 mm petri dish and oriented ventral side up to the imaging plane. Embedded embryos were transferred to an imaging chamber (RC-29; Harvard Instruments, USA) with a heated platform (PH-6D; Harvard Instruments). Temperature was maintained at 28°C by a temperature controller (TC-344B; Harvard Instruments). Images were obtained using an epifluorescence upright microscope (BX51WI; Olympus) and focusing module (BXFM; Olympus) with a 40X water immersion objective (LUMPLFLN 40XW; Olympus) and magnification changer (U-CA; Olympus). Fluorescence was excited using a blue light-emitting diode (CBT-90; Luminus, USA) passed through a 460±5 nm bandpass filter (HQ460/10X; Chroma, USA). Fluorescence was collected with a 482 nm dichroic mirror (FF482-Di01; Semrock, USA). To obtain simultaneous images of FRET donor and acceptor signals, collected light was passed into an image splitter (OptoSplit II; Cairn Research, USA), split with a 552 nm dichroic mirror (FF552-Di02; Semrock), and passed through either a 500±30 nm (HQ500/60m-2p; Chroma) or 600±37.5 nm (HQ600/75m; Chroma) bandpass filter. Filtered emission was projected to two halves of a 16-bit, 128×128, 24 mm^2^ pixels, cooled electron multiplying charge-coupled device camera (Cascade: 128+; Photometrics, USA) and collected at 52Hz with a ∼19 ms exposure time. Images were processed and analyzed using custom routines in MATLAB (R2011b; MathWorks, USA). The ratio of FRET donor and acceptor signals was taken and spatially filtered using the pixelwise adaptive, linear, noise-removal Wiener method (‘wiener2’) with a 3×3 pixel window. The atrium was manually segmented and each pixel signal normalized through time. Activation time was measured as the point at which the rate of voltage upstroke was maximal.

### Electrophysiology

Micropipettes for electrocardiograph (ECG) measurement on whole zebrafish larvae were prepared by pulling fire-polished borosilicate glass capillaries (World Precision Instruments) using the Flaming/brown micropipette puller P-1000 (Sutter Instrument). The zebrafish larvae were mounted (laterally) in 1% low melting agarose in a glass dish and submerged in external buffer: 1x egg water (0.6g/L sea salt in reverse osmosis purified water). The micropipette was filled with internal buffer (174mM NaCl, 2.1mM KCL, 1.2mM MgSO4.7H20, 1.8mM Ca(NO3)2.4H2O, 15mM HEPES, pH 7.2) and the tip was positioned right above the pericardial region of the zebrafish heart. The electrical signals from the zebrafish heart received were recorded by pCLAMP 10 software (Molecular Devices) after amplification via Multiclamp 700B amplifier (Molecular Devices) and digitization through Axon Digidata 1440A digitizer (Molecular Devices). Data were analysed with Clampfit 10 software (Molecular Devices).

### Whole mount *in situ* hybridization

For antisense probes generation, total RNA from 72 hpf embryos was extracted and reverse transcribed into cDNA with SuperScript IV Reverse Transcriptase (Thermo Fisher Scientific, USA). Obtained cDNA was used as a template for PCR. Purified PCR products were used as a template for *in vitro* transcription from the T7 promoter. Primers used are listed in S1 Table or reported previously [51]. Whole mount *in situ* hybridization (WISH) was performed as previously described, with minor adjustments [120]. Zebrafish embryos were grown in egg water containing PTU and fixed overnight at desired developmental stage in 4% paraformaldehyde in 1x PBS (PFA/PBS). After sequential washes with 1x PBT (50 ml 1x PBS + 250 µl 20% Tween-20), embryos were digested for either 30 min (48 hpf) or 50 min (72 hpf) with 10 µg/ml proteinase K (Roche), washed with 1x PBT, and fixed again for 1 hour. PFA/PBS solution was discarded, and embryos were pre-hybridized overnight at 68°C in a hybridization buffer. Subsequently, diluted and denatured probes were added to the pre-hybridized embryos followed by overnight incubation (68°C) in a water bath. Post-hybridization washes were performed in increasing concentration of 2xSSC in the hybridization buffer. To reduce nonspecific signal, commercial blocking reagent (Roche) was used. Signal was visualized by overnight incubation with 1:5000 anti-DIG-AP antibody (Roche) at 4°C followed by washing and addition of NBT and BCIP staining solution. After the staining was fully developed, staining solution was washed away and embryos were fixed in 4% PFA in 1x PBS. For whole mount *in situ* imaging, embryos were mounted in glycerol and imaged on Nikon SMZ25 microscope. For each probe, WISH experiments were performed on at least 20 embryos obtained from at least 3 different breeding pairs.

## Acknowledgements

We are grateful to the zebrafish core facility of the IIMCB Warsaw for excellent fish care. We thank Dr. Y. Okamura for the Tol2-*myl7:mermaid* construct and Dr. Natascia Tiso for kindly sharing the *Tg(7x TCF-Xla.Sia:NLS-mCherry)* transgenic line. We thank W. Rybski and Y. Siddiqui for technical help, Drs. D. Stainier, T. Braun, L. Solnica-Krezel, R. Minhas for providing critical advice and fruitful discussions.

## Supporting information

**S1 Figure.**
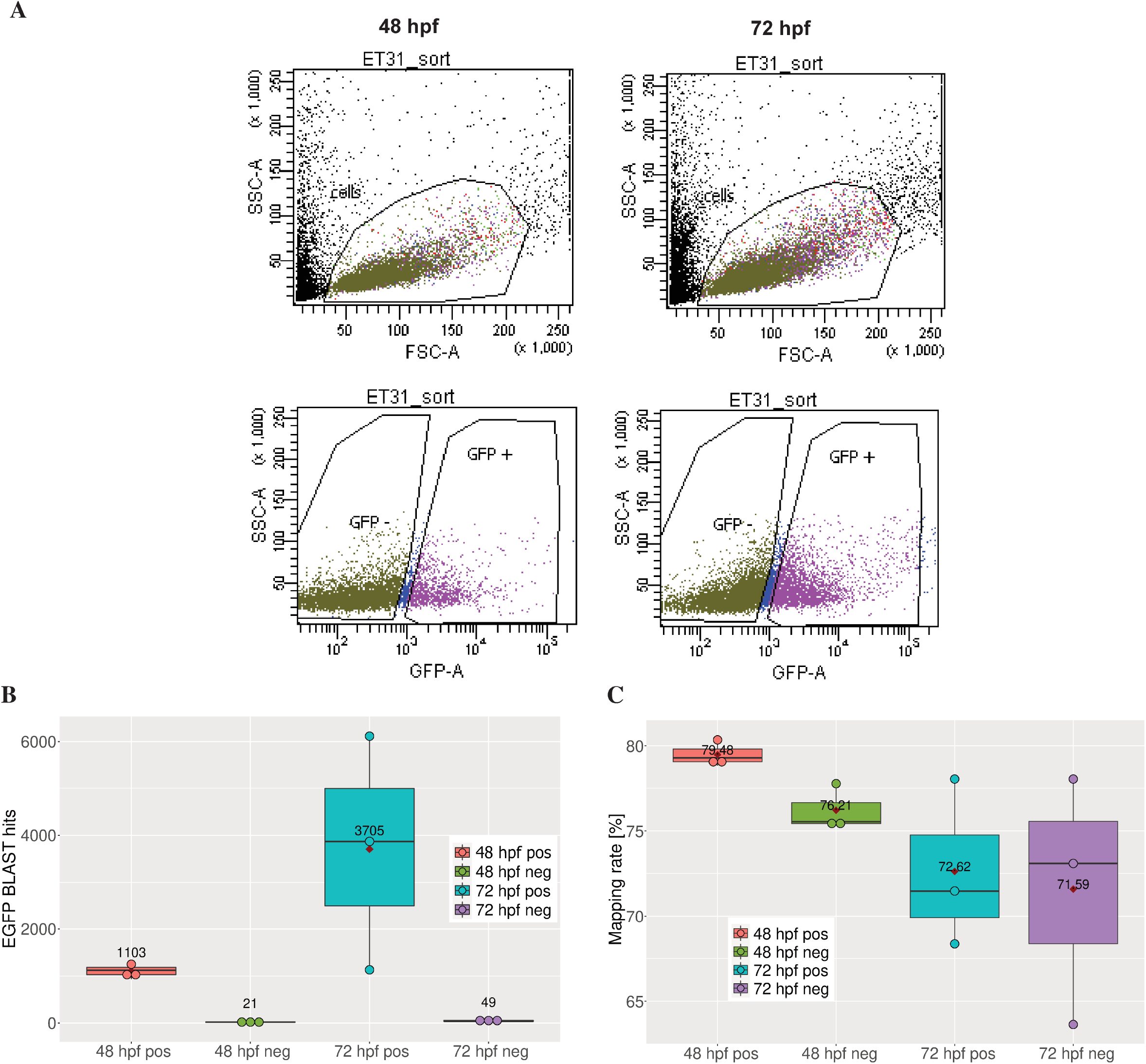
Sample quality measures. (A) Analysis of FACS sorting of cardiac cells from the *sqet31Et* transgenic line. Forward scatter (FSC-A) vs side scatter (SSC-A) was used to find viable, single cell events; SSC-A vs GFP-A was applied to identify fluorescent cells. (B) Box plot showing distribution of EGFP transcript levels as a result of BLAST search against EGFP sequence in three different replicates of each sample type. Note the significant enrichment of EGFP expression in EGFP-positive samples as compared to negative, which is reflected by the fold change 52.5 and 75.6 for 48 hpf and 72 hpf, respectively. Mappability of RNA-seq reads in three replicas of each sample type. (C) Box plot of sequencing reads mapping rates in each sample type. Average mapping rate for all the samples was around 75%.

**S2 Figure.**
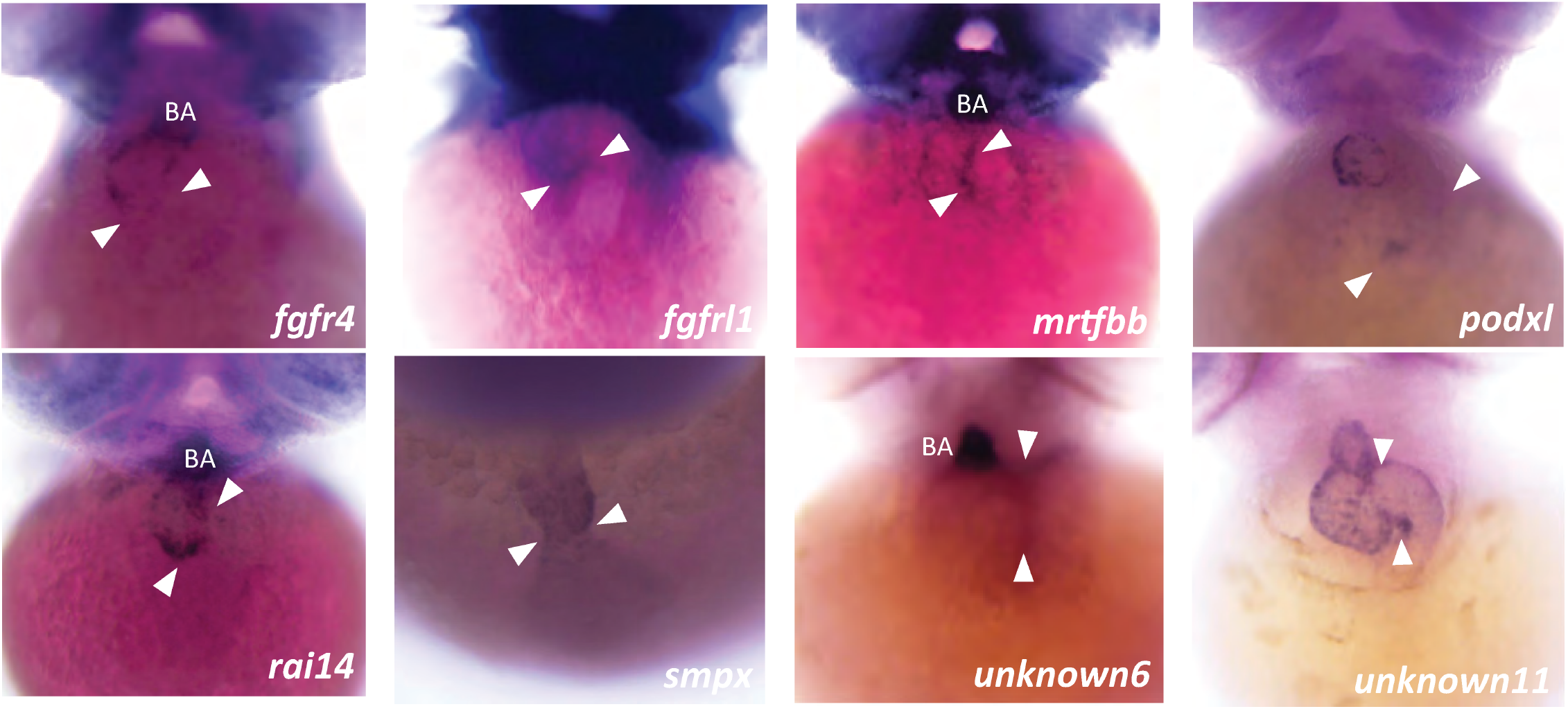
Expression validation of AVC-enriched genes by whole mount *in situ* hybridization at 72 hpf, except for *smpx* (48 hpf). Images show the frontal view of the embryos. The AVC region is indicated by white arrowheads. BA – bulbus arteriosus.

**S3 Figure.**
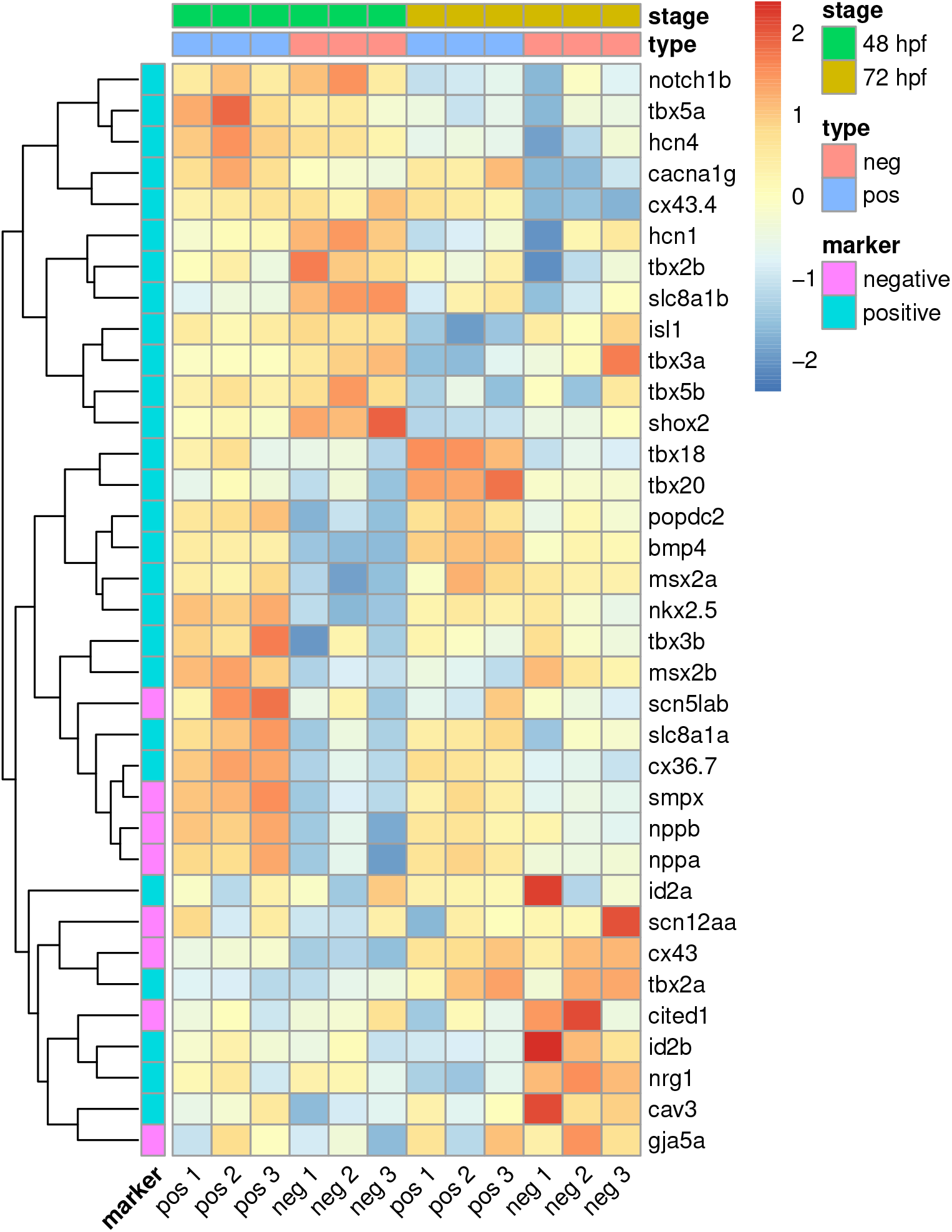
Overview of known gene signatures associated with the atrioventricular node and pacemaker. In general, expressed as a normalized expression value (regularized log [rld]). Both positive and negative markers (working myocardium) were shown.

**S4 Figure.**
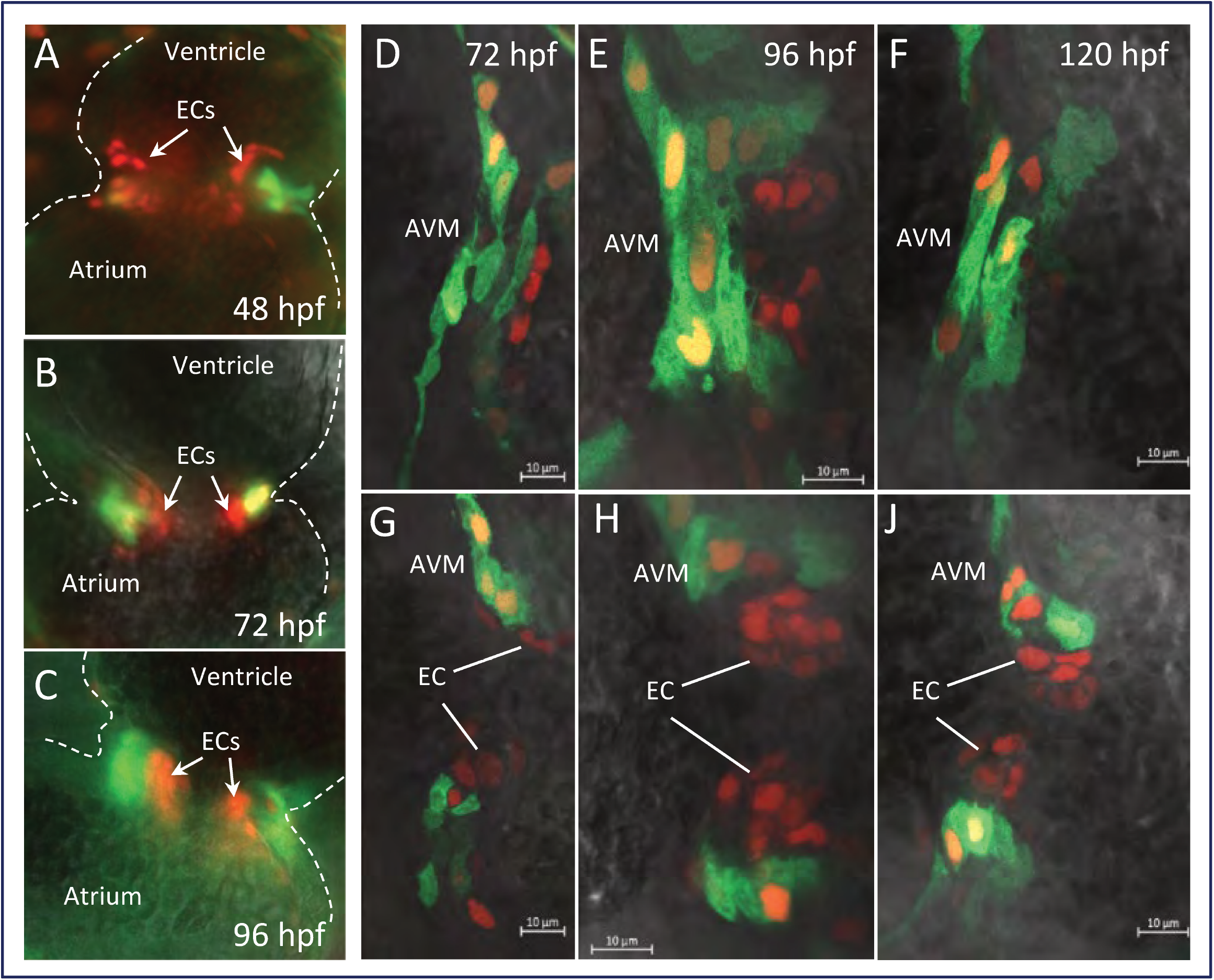
Wnt signaling activity in different layers of the AVC. Double transgenic line *sqet31Et* x *Tg(7xTCFXla.Siam:nlsmCherry)* were generated in order to visualize Wnt signaling activity (represented by mCherry expression) in the AVC region. (A-C) Fluorescent image showing Wnt signaling activity in cells of the endocardial cushion (ECs). (C - H) confocal image of the AVC at the surface (C-E) and lumen (F-H) reveals overlap of mCherry expression with that of EGFP in the *sqet31Et* line, indicating additional Wnt signaling activity in the AV myocardium (AVM).

**S5 Figure.**
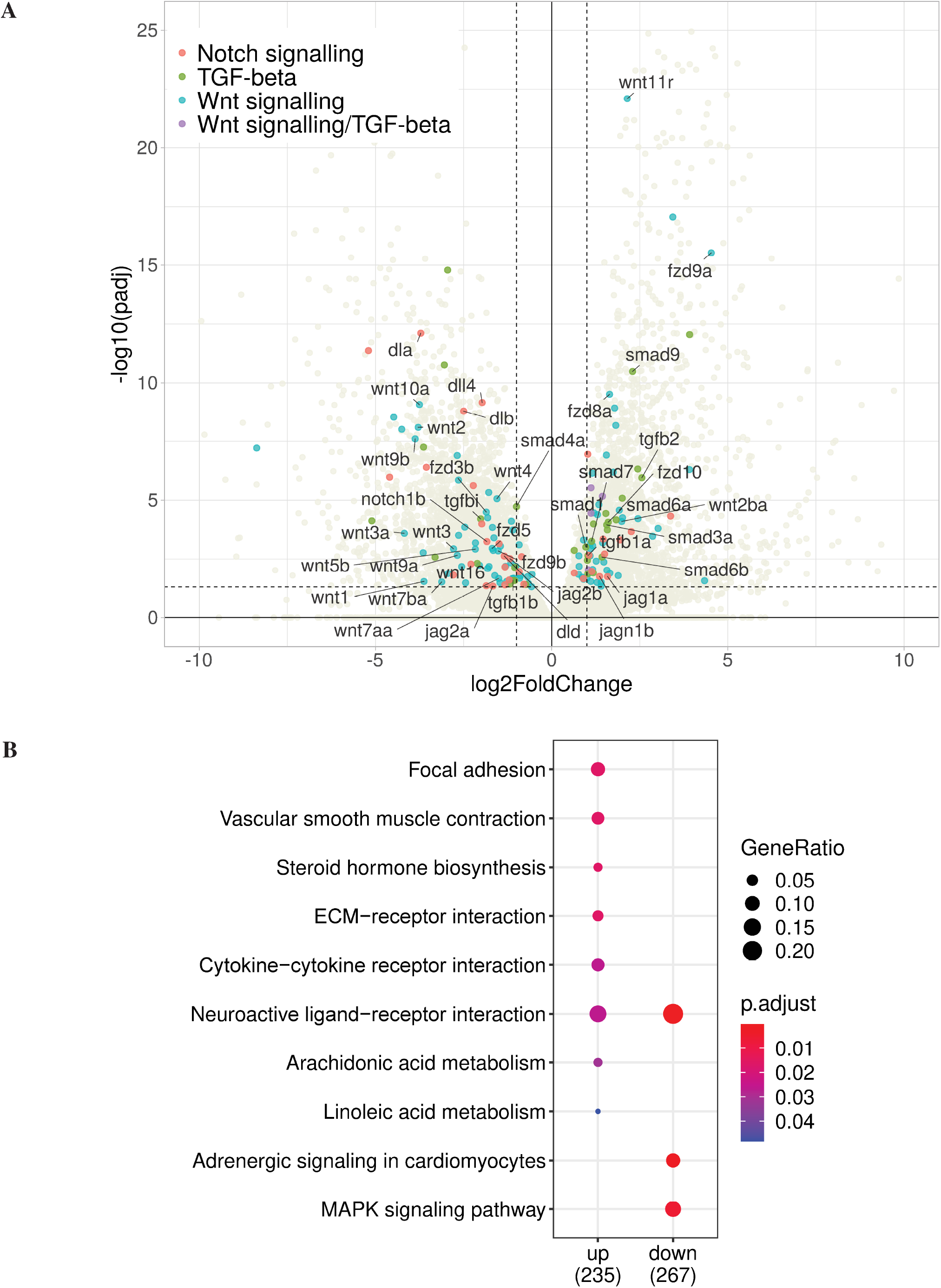
Developmental signaling pathway genes with dynamic expression between 72 hpf and 48 hpf. (A) Volcano plot representing the expression of various components of Notch, TGF-β and Wnt signalling pathways in the AVC myocardium (padj < 0.05, -1 < log2FoldChange > 1). (B) Overrepresented Gene Ontology terms at 72 hpf as compared to 48 hpf.

**S1 Table. List of primers for *in situ* probes.**

**S2 Table. Master list of GFP+ vs - differentially expressed genes at both stages. S3 Table. GO enrichment analysis of enriched genes at both stages.**

**S4 Table. Common AVC gene signatures between 48 hpf and 72 hpf.**

**S5 Table. Connexins enriched in the GFP+ population at both developmental stages.**

**S6 Table. Genes associated with atrioventricular node or pacemaker development and function.**

**S7 Table. AVC and SAR intersection.**

**S8 Table. Genes involved in the EMT process enriched in the GFP+ compared to GFP-cell population.**

**S9 Table. Master list of GFP+ differentially expressed genes between 72 hpf vs 48 hpf. S10 Table. GO analysis of differentially expressed genes at 72 hpf vs 48 hpf.**

**S11 Table. List of significant gene members of three signaling pathways.**

**S12 Table. Results of ClinVar search database in terms of GWAS study related to the cardiac conduction system.**

**S13 Table. Results of ClinVar search database in terms of GWAS study related to the valve.**

**S14 Table. Results of ClinVar search database in terms of GWAS study related to the heart septal defects.**

**S15 Table. Bulbus arteriosus genes enriched in GFP+ cells at 72 hpf.**

